# The endosomal sorting adaptor HD-PTP is required for ephrin-B:EphB signalling in cell collapse and motor axon guidance

**DOI:** 10.1101/386631

**Authors:** Sylvie Lahaie, Daniel Morales, Halil Bagci, Noumeira Hamoud, Charles-Etienne Castonguay, Jalal M. Kazan, Guillaume Desrochers, Avihu Klar, Anne-Claude Gingras, Arnim Pause, Jean-François Côté, Artur Kania

## Abstract

The signalling output of many transmembrane receptors that mediate cell-cell communication is restricted by the endosomal sorting complex required for transport (ESCRT), but the impact of this machinery on Eph tyrosine kinase receptor function is unknown. We identified the ESCRT-associated adaptor protein HD-PTP as part of an EphB2 BioID interactome, and confirmed this association using co-immunoprecipitation. Although HD-PTP loss does not change EphB2 expression, it attenuates the ephrin-B2:EphB2 signalling-induced collapse of cultured cells and axonal growth cones, and results in aberrant guidance of chick spinal motor neuron axons *in vivo* HD-PTP depletion abrogates ligand-induced EphB2 clustering, and EphB2 and Src family kinase activation. HD-PTP deficiency also accelerates ligand-induced EphB2 degradation, contrasting the phenotypes reported for other cell surface receptors. Our results link Eph signalling to the ESCRT machinery and demonstrate a role for HD-PTP in the earliest steps of ephrin-B:EphB signalling, as well as in obstructing premature receptor depletion.

## Introduction

Cell-cell contact-dependent signalling underlies many diverse biological processes such as tissue boundary formation, synaptic plasticity, axon guidance, and tumorigenesis. The relatively small family of Eph receptor tyrosine kinases plays a major role in all of these, but the molecular pathways that restrict Eph signalling with impressive spatiotemporal precision are still being unravelled (Kania & Klein, 2016). The highly-conserved endosomal sorting complex required for transport (ESCRT) modulates the signalling of many classes of cell surface receptors through their internalisation, lysosomal degradation or recycling (Raiborg & Stenmark, 2009). Intriguingly, despite ESCRT’s nearly universal involvement in transmembrane receptor function, its role in Eph signalling remains unexplored.

Eph receptor A and B subfamilies are defined by their ephrin ligands’ linkage to the cell membrane via a GPI anchor or a transmembrane domain, respectively. Forward signalling evoked by ephrin binding to the Eph ligand binding domain (ephrin:Eph) typically results in a rapid and restricted actin cytoskeleton collapse in the Eph expressing cell, underlying cell-cell repulsion at tissue boundaries, cancer cell invasion, and dendritic spine plasticity (Pasquale, 2010; Klein, 2012). Arguably, the best understood *in vivo* ephrin:Eph signalling events are those directing axonal growth cones; for example, ephrin-B ligands expressed by the vertebrate dorsal limb mesenchyme, repel EphB-expressing spinal motor neuron axons and direct them to their muscle targets in the ventral limb (Luria *et al.*, 2008). At the molecular level, one early critical event in ephrin:Eph signalling is the formation of large Eph multimer arrays upon ephrin binding (Himanen *et al.*, 2010; Seiradake *et al.*, 2010). The induction of Eph clusters is sufficient to induce cytoskeletal collapse (Egea *et al.*, 2005), and their size and composition determine the strength of this response (Schaupp *et al.*, 2014). Besides ephrin-Eph contacts, clustering is driven by Eph-Eph interactions via Eph extracellular cysteine-rich domains (Himanen *et al.*, 2010), intracellular SAM domains (Thanos *et al.*, 1999) and, possibly, PDZ domain-containing intracellular adaptor proteins (Torres *et al.*, 1998). Eph clustering enables the phosphorylation of juxtamembrane tyrosines which is required for the activation of the Eph kinase domain (Egea *et al.*, 2005; Binns *et al.*, 2000), resulting in the recruitment of intracellular effectors including Src family kinases, linking receptor activation to the actin cytoskeleton (Ellis *et al.*, 1996; Zisch *et al.*, 1998). Despite the critical importance of receptor clustering in the initiation of the Eph signalling cascade, the factors that control it remain virtually unknown.

The endosomal internalisation of ephrin:Eph receptor complexes is required for their normal signalling (Marston *et al.*, 2003; Zimmer *et al.*, 2003, Cowan *et al.*, 2005), and eventually leads to dephosphorylation of juxtamembrane tyrosines (Shintani *et al.*, 2006), ubiquitylation of the Eph cytoplasmic tail (Okumura *et al.*, 2017) and Eph recycling or degradation (Sabet *et al.*, 2015). It is unknown whether the fate of internalised Eph receptors depends on the ESCRT machinery, which detects ubiquitylated receptors and transfers them between specialised vesicles where they are subject to deubiquitylation, and sorting to the lysosome (Raiborg & Stenmark, 2009; Szymanska *et al.*, 2018). Among the regulators of this progression is the Bro1 domain-containing cytosolic protein, His-domain-containing protein tyrosine phosphatase (HD-PTP, also known as PTPN23 and Myopic), which brings ESCRT proteins directly in contact with the UBPY deubiquitylase (Ali *et al.*, 2013; Gahloth *et al.*, 2017). HD-PTP loss leads to impaired sorting of internalised receptors and their aberrant accumulation in endosomes (Doyotte *et al.*, 2008; Kharitidi *et al.*, 2015). Mice heterozygous for *Ptpn23*, the gene encoding HD-PTP, are predisposed to various tumours (Manteghi *et al.*, 2016), a phenotype commonly associated with excessive activation of morphogen and growth factor receptors. HD-PTP has not been studied in the context of Eph signalling, and more generally, the only evidence linking Eph signalling to the ESRCT machinery is the observation that EphB2 can associate with ESCRT proteins in the context of exosome biogenesis (Gong *et al.*, 2016).

To study the proteomic environment of activated Eph receptors and its relation to Eph clustering and endocytic sorting, we performed a proximity-dependent biotin identification experiment in cells expressing EphB2, exposed to ephrin-B2. Among the identified EphB2-proximal proteins we found HD-PTP, which can interact with EphB2 in a ligand-dependent manner, and is required for EphB2 signalling in the context of cell and growth cone collapse, as well as in the guidance of spinal motor neuron axons *in vivo*. Our experiments argue that HD-PTP functions in the earliest steps of the Eph signalling cascade: the formation of EphB2 clusters and Src family kinase phosphorylation in response to ephrin-B2 stimulation. HD-PTP also acts at a later step in the Eph signalling pathway, although in contrast to its role in ESCRT processing of other receptors, it acts as a negative regulator of EphB2 degradation. Altogether, these results are the first to establish a functional link between Eph signalling and ESCRT accessory proteins, revealing their novel role in promoting cell surface receptor signalling.

## Results

### A BioID survey of the ephrin-B2-induced EphB2 interactome

To identify new proteins potentially involved in EphB2 receptor activation and its processing, we used proximity-dependent biotin identification (BioID; Roux *et al.*, 2012). We generated a Flp-In T-REx HEK293 cell line with inducible expression of EphB2-BirA*-FLAG (EphB2-OE HEK), where the BirA* biotin ligase was fused to the C-terminus of EphB2, allowing us to identify proteins in close proximity to EphB2 during eB2-induced forward signalling. We stimulated these cells with either clustered ephrin-B2-Fc (eB2), Fc or media (“no ligand”) for 6 h, followed by lysis, streptavidin pull-down and mass spectrometry (MS; *n* = 4 per condition; Fig. 1A, B). MS data were filtered using Significant Analysis of INTeractome (SAINT; Teo *et al.*, 2014), with BirA*-FLAG-EGFP and empty vector HEK293 MS datasets as a controls, yielding prey peptides with a Bayesian false discovery rate (BFDR) score ≤ 0.01 (Supplementary Table 5). We analysed eB2 and Fc SAINT datasets by calculating each prey’s WD-Score, a measure of hit specificity (Knight *et al.*, 2017). Differences between average spectral counts and WD-scores in eB2 or Fc conditions were found for many preys (Fig. 1C; Supplementary Table 5).

**Figure 1.**
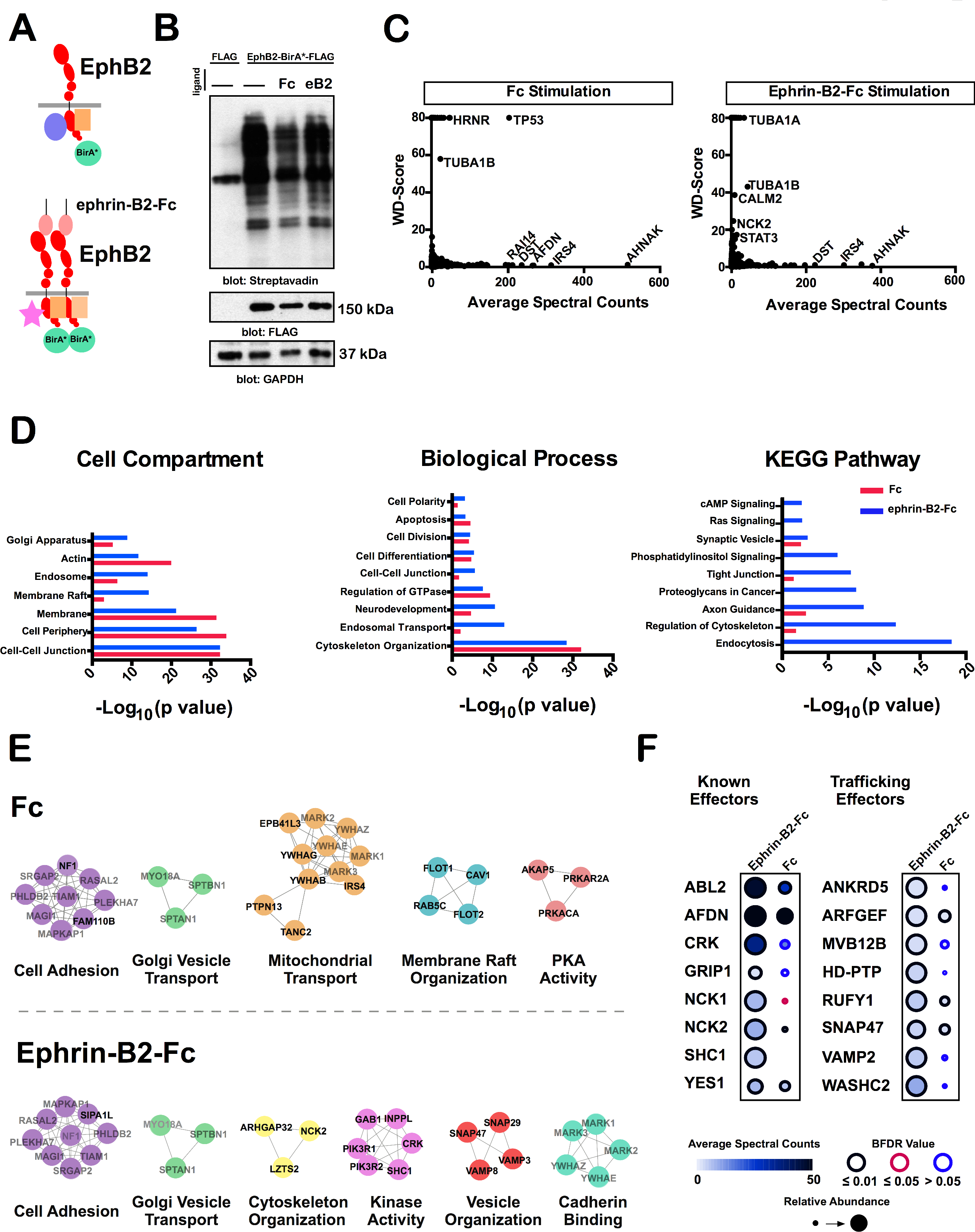
A BioID screen for ligand-stimulated EphB2-proximal proteins. A) Schematic of the BioID experiment: a FLAG-tagged biotin ligase, BirA*, was fused to the C-terminus of EphB2 and stably expressed in HEK293 cells. Depending on the presence of ephrin-B2 ligands, different proteins are recruited to the vicinity of EphB2. B) Western blot of biotinylation in protein lysates from our mass spec samples of HEK293 cells expressing EphB2-BirA*-FLAG (lane 2, no ligand; 3, 1.5 μg/mL Fc; and 4, 1.5 μg/mL pre-clustered ephrin-B2-Fc) or FLAG alone (lane 1). (*n* = 2). C) Plots of WD-Score *vs*. average spectral count in eB2-treated samples and Fc only treated controls. Each point represents a protein identified by MS (*n* = 4). D) Gene ontology and KEGG terms associated with the proteins identified in the eB2 WD-score and Fc WD-score analysis. Blue bars represent proteins enriched in the eB2-treated samples and red bars represent proteins enriched in the Fc-treated samples. E) Interactome webs of proteins identified in the eB2 or Fc WD-score analysis generated by Cytoscape and clustered with MCluster, divided by gene ontology term. F) Dot plots for known EphB2 effector or trafficking-related preys in eB2 and Fc conditions. Spectral count is illustrated by fill shade, relative abundance of the protein compared to the EGFP-BirA*-FLAG condition shown by circle size, and outer circle color represents BFDR value when compared to EGFP-BirA*-FLAG MS SAINT analysis. kDa: kilodalton; eB2: ephrin-B2-Fc; BFDR: Bayesian false discovery rate.

Next, we used g:Profiler to perform functional annotations of the WD-score analysis of eB2 and Fc conditions, which showed an enrichment of proteins associated with endosomal trafficking and neurodevelopmental biological processes in the eB2-stimulated profile (Reimand *et al.*, 2016; Fig. 1D). Using the Cytoscape database (Shannon *et al.*, 2003) and the Markov Cluster (MCL) tool (Enright *et al.*, 2002), we generated two interactome maps using Fc and eB2 WD-Score analysis (Fig. 1E). As expected, eB2-stimulated protein clusters include known EphB2 forward signalling functions such as cytoskeleton organization, kinase activity, and vesicle organization.

Based on these broad visualisations of our MS data, we examined individual preys more specifically, comparing their average spectral counts, relative abundance, and BFDR score between the eB2 and Fc treatments (Fig. 1F). Several known EphB2-binding proteins were enriched upon eB2 stimulation, such as Abelson kinase (ABL2; Yu *et al.*, 2001) and Nck adaptor proteins (NCK1 & NCK2; Stein *et al.*, 1998), suggesting that our BioID analysis sampled the protein environment of active EphB2 forward signalling. Among the trafficking proteins that were enriched by eB2 stimulation, we found HD-PTP, a known ESCRT adaptor protein with trafficking functions but without previous evidence of involvement in Eph signalling (Raiborg & Stenmark, 2009).

### HD-PTP and EphB2 expression and localisation are linked

To determine whether EphB2 and HD-PTP can form a complex, we performed co-immunoprecipitation (co-IP) assays in the EphB2 HEK cell line transfected with an HD-PTP-HA expression plasmid. Following EphB2-BirA*-FLAG induction, the cells were treated with ephrin-B2-Fc, Fc, or media (Fig. 2A). The EphB2-FLAG-directed pull-down showed a stronger anti-HA band following eB2 stimulation compared to Fc or media conditions (Fig. 2B, C; *n* = 4; *p* = 0.0017) suggesting that HD-PTP can form a complex with EphB2 and the efficacy of this effect is increased by eB2.

**Figure 2.**
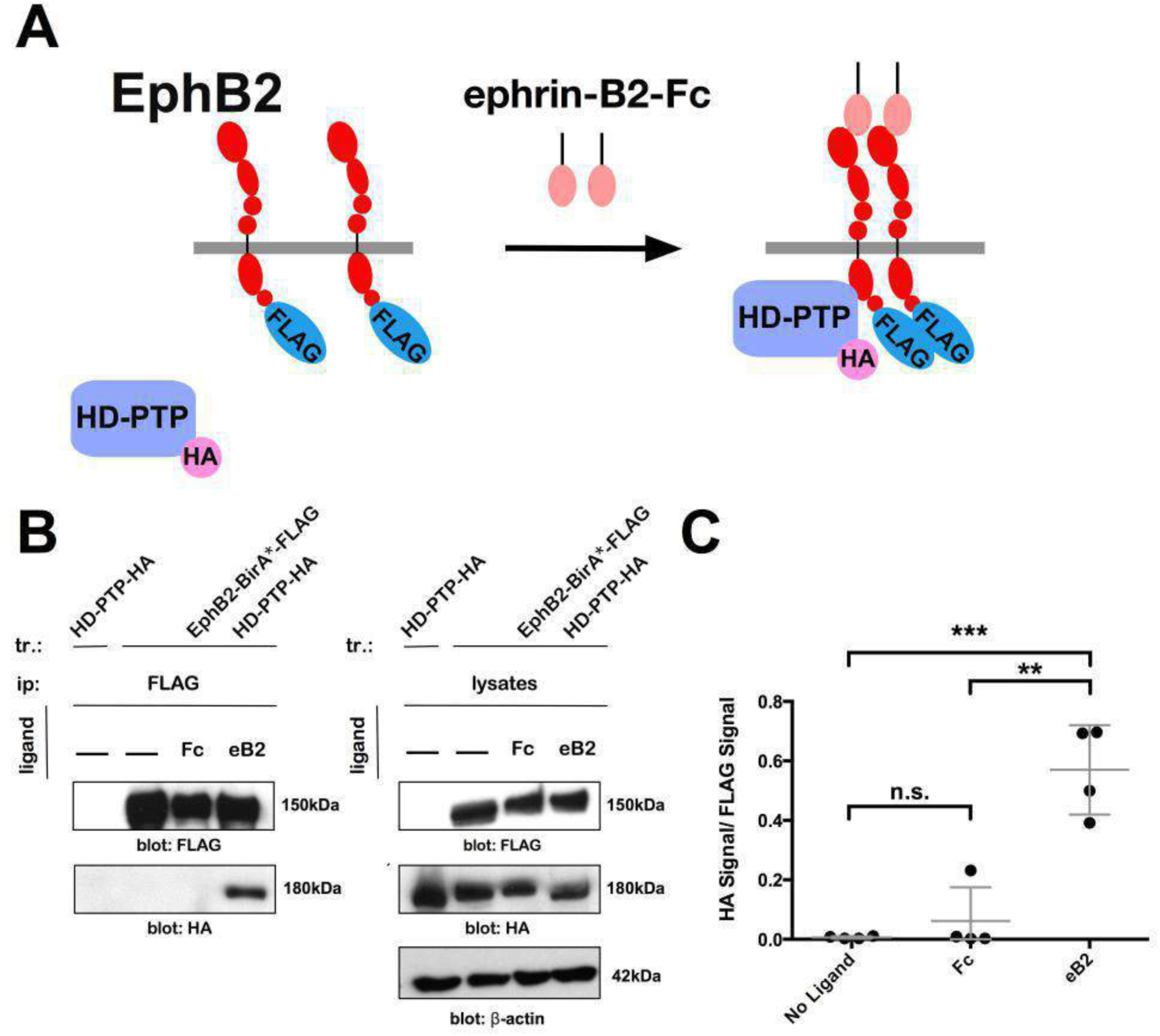
EphB2 and HD-PTP can form a ligand-dependent complex. A) Schematic depicting ligand-dependent HD-PTP complex-formation with EphB2 and location of FLAG and HA tags. B) Representative blot of a co-immunoprecipitation experiment performed in HEK293 cells transfected with HD-PTP-HA and expressing either EphB2-BirA*-FLAG or FLAG alone. Blotting for HA shows enhanced pull-down of HD-PTP with EphB2 in cells that have been stimulated for 15’ with 1.5 μg/mL eB2. C) Quantification of HA signal in co-immunoprecipitation experiments, normalised to FLAG. More signal is detected in eB2-treated cells than in Fc or no ligand controls, *p* = 0.0017; one-way ANOVA followed by Student’s *t*-tests corrected for multiple comparisons. Mean of four independent experiments. Values are plotted as mean ± SD. All values can be found in Supplementary Table 4. kDa: kilodalton; eB2: ephrin-B2-Fc; *** *p* < 0.001; ** *p* < 0.01; n.s.: not significant.

Next, we examined whether HD-PTP and EphB2 expression and localisation are linked. In a tetracycline-inducible FLAG Flp-In T-REx HeLa cell line (Control HeLa; nomenclature in Supplementary Table 3), we found a significant correlation between the expression levels of both proteins in individual cells (Fig. 3D; *n* = 82 cells; *R^2^* = 0.218; *p* < 0.0001). We also compared this relationship in a HeLa cell line with HD-PTP levels reduced through expression of an *HD-PTP* short hairpin RNA (Kharitidi *et al.*, 2015; HD-PTP^shRNA^), in HeLa cells with an empty shRNA viral vector (Control^shRNA^ HeLa) and in Control^shRNA^ HeLa cells transfected with an *HD-PTP* expression plasmid (HD-PTP-OE; Fig. 3A, C). Neither HD-PTP loss or overexpression produced a change in EphB2 signal levels (Fig. 3A, B; *n* = 3; *p* = 0.9992). In contrast, HeLa cells with increased EphB2 expression levels (EphB2-OE HeLa) had a significantly increased HD-PTP expression compared to Control HeLa cells (Fig. 3E-I), without similar effects on the levels of another intracellular protein, BEN (Sup. Fig. 3A, B; *n* = 3; *p* = 0.3695). Finally, in EphB2-OE HeLa cells approximately 80% of HD-PTP signal localised to EphB2^+^ puncta, a significant difference from controls (Fig. 3J, K). Together, these data argue that EphB2 and HD-PTP can form a molecular complex, and their expression and localisation are linked.

**Figure 3.**
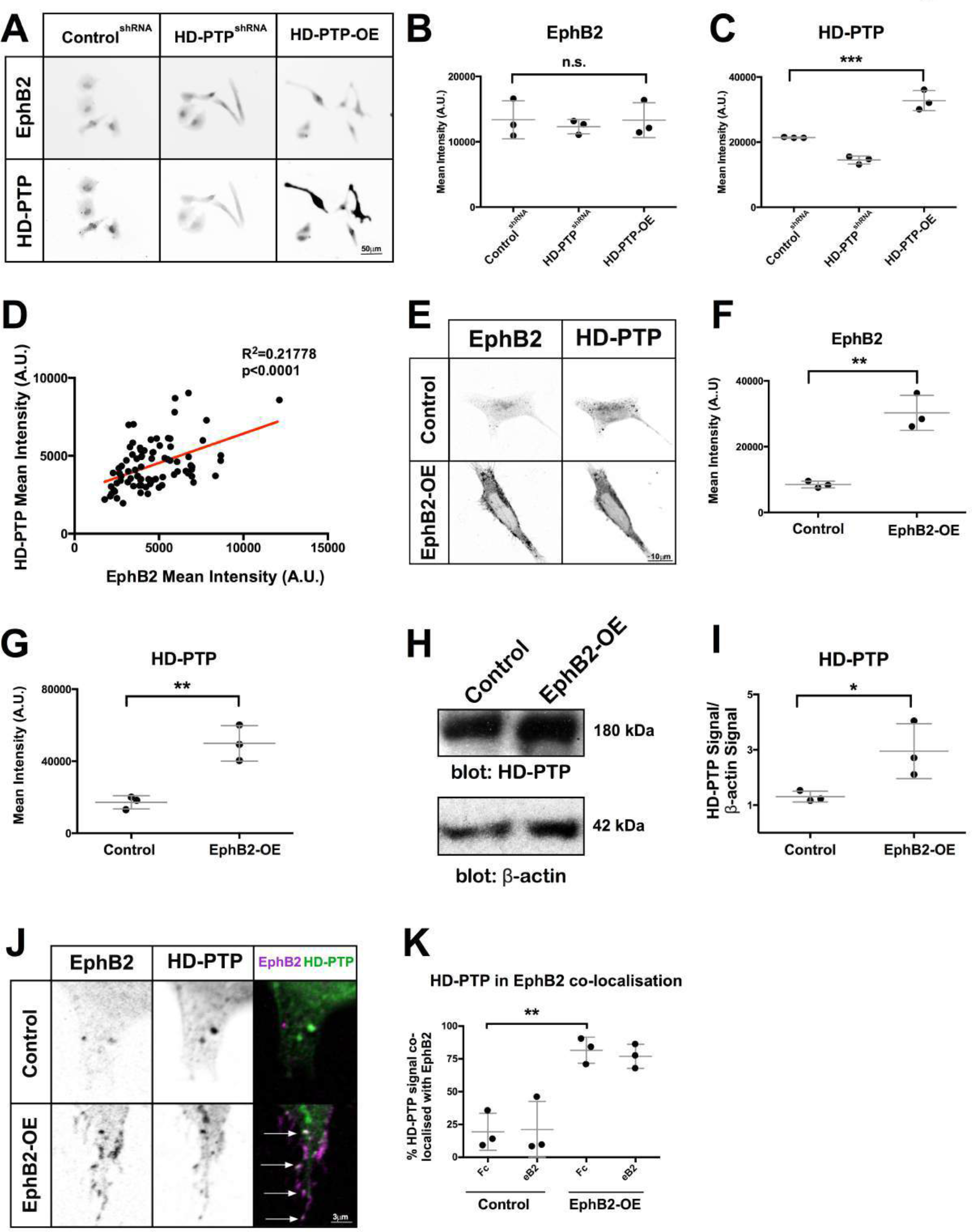
HD-PTP and EphB2 expression and localisation are linked in HeLa cells. A) Representative examples of EphB2 and HD-PTP expression visualised by immunohistochemistry in Control^shRNA^, HD-PTP^shRA^, and HD-PTP-OE HeLa cells. B) Quantification of anti-EphB2 mean pixel intensity in Control^shRNA^, HD-PTP^shRNA^, and HD-PTP-OE HeLa cells. Expression of EphB2 is not changed in any condition (*n* = 3; *p* = 0.9992; one-way ANOVA followed by corrected Student’s *t*-tests). C) Quantification of anti-HD-PTP mean pixel intensity in Control^shRNA^, HD-PTP^shRNA^, and HD-PTP-OE HeLa cells. HD-PTP signal is reduced by 32% in HD-PTP^shRNA^ versus Control^shRNA^, and increased by 53% in HD-PTP-OE versus Control^shRNA^ (*n* = 3, 60–80 cells/*n*; one-way ANOVA followed by corrected Student’s *t*-tests). D) Scatter plot of HD-PTP *vs*. EphB2 signal intensity in Control HeLa cells. Each data point represents a cell from 3 experiments. There is a significant correlation between the mean intensity levels of the two proteins (*n* = 3, 20–30 cells/*n*; *R^2^* = 0.218; Y=0.3765*X+2658; *p* < 0.0001; simple linear regression and correlation analysis). E) Representative examples of EphB2 and HD-PTP expression visualised by immunohistochemistry in Control and EphB2-OE HeLa cells. F) Quantification of EphB2 mean pixel intensity signal shows a three-fold increase in EphB2-OE HeLa compared to Control HeLa (*n* = 3, 60–80 cells/*n*; Student’s *t*-test). G) Quantification of HD-PTP mean signal pixel intensity shows a two-fold increase in EphB2-OE HeLa compared to Control HeLa (*n* = 3, 60–80 cells/*n*; Student’s *t*-test). H) Western blot detection of HD-PTP and β-actin in lysates from Control and EphB2-OE HeLa cells. I) Quantification of HD-PTP Western blot signal normalised to β-actin shows increased HD-PTP levels in EphB2-OE HeLa cell lysate compared to Control HeLa (*n* = 3; Student’s *t*-test). J) Representative images of EphB2-OE and Control HeLa cells stained with anti-EphB2 and anti-HD-PTP antibodies showing co-localisation of HD-PTP and EphB2. K) Quantification of HD-PTP signal localisation in EphB2-positive domains in HeLa cells. HD-PTP is preferentially found in EphB2-containing puncta in EphB2-OE HeLa cells compared to Control HeLa cells (*n* = 3, 10–12 cells/*n*; Student’s *t*-test). Values are plotted as mean ± SD. All values can be found in Supplementary Table 4. kDa: kilodalton; eB2: ephrin-B2-Fc; *** *p* < 0.001; ** *p* < 0.01; * *p* < 0.05; n.s.: not significant. Scale bars: A) 50 μm, E) 10 μm, J) 3 μm. Inverted grayscale fluorescent images except for dual colour images in J.

### HD-PTP is required for ephrin-B2-induced cell collapse

We next assessed whether HD-PTP functions in ephrin-B2:EphB2 signalling, which in many contexts causes a destabilisation of the cytoskeleton. Using immunohistochemistry, we observed that in Control HeLa cells, EphB2 signal intensity was inversely correlated with cell size (Fig. 4A; *p* < 0.0001; R^2^ = 0.31), suggesting that, similar to ephrin-A:EphA signalling-induced HeLa cell collapse, EphB2 may reduce HeLa cell size by responding to endogenously expressed ephrin-Bs (Seiradake *et al.*, 2013; Thul *et al.*, 2017). Control HeLa cells treated with eB2 showed a ~20% reduction in cell area compared to cells treated with Fc (Fig. 4B, C; *n* = 4; *p* = 0.0046), while EphB2-OE HeLa cells treated with eB2 were ~50% smaller than those incubated with Fc (Fig. 4B, C; *n* = 8; *p* = 0.0004). Together with the observation that increasing concentrations of eB2 cause greater reduction in EphB2 HeLa cell size (Fig. 4D; *n* = 4), our data suggest that eB2:EphB2 signalling causes HeLa cell collapse.

**Figure 4.**
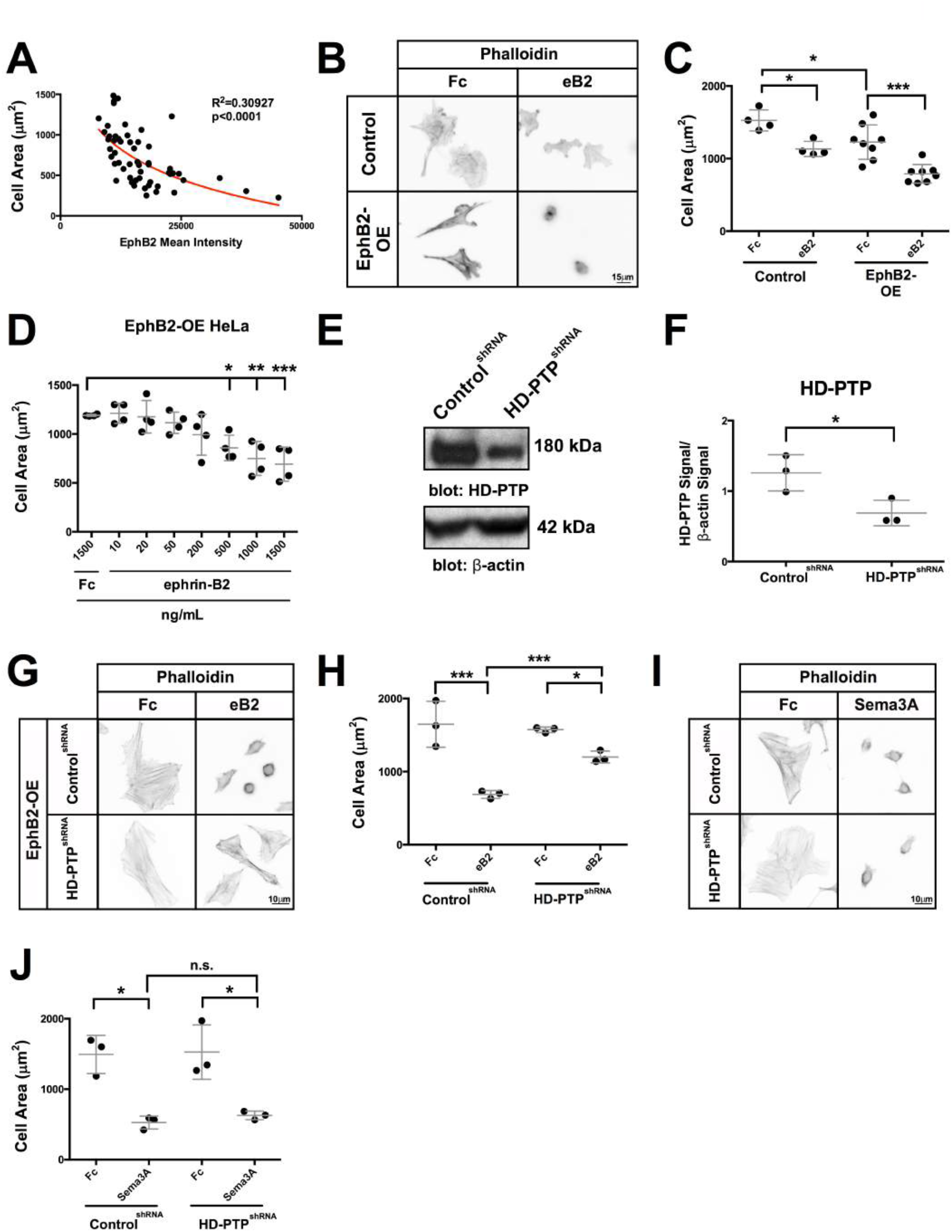
HD-PTP is required for ephrin-B2-induced cell collapse. A) Scatter plot of cell area *vs*. EphB2 mean pixel intensity shows a strong negative correlation in Control HeLa cells (*n* = 67; *p* < 0.0001; R^2^ = 0.31; the curve fit equation is Y=541.7(-4.842e^-0.005^ *X). B) Representative images of Control HeLa and EphB2-OE HeLa cells, stimulated 15’ with 1.5 μg/mL eB2 or Fc and stained with phalloidin conjugated with Alexa Fluoro 568, revealing the actin cytoskeleton. C) Quantification of cell size of HeLa cells treated with eB2 or Fc. Ligand-induced cell collapse was evident in Control cells, and in EphB2-OE HeLa cells (*n* = 4, 60–80 cells/*n* in Control, *p* = 0.0046; *n* = 8, 60–80 cells/n in EphB2-OE, *p* = 0.0004; one-way ANOVA followed by corrected Student’s *t*-tests). D) EphB2-OE HeLa cell size following treatment with increasing concentrations of eB2 for 15’ (*n* = 4, 60–80 cells/*n*; one-way ANOVA followed by corrected Student’s *t*-tests). E) Representative Western blot of HD-PTP and β-actin in lysates of HeLa cells stably expressing Control^shRNA^ and HD-PTP^shRNA^. F) Quantification of HD-PTP Western blot signal normalised to β-actin shows decreased HD-PTP protein levels in HD-PTP^shRNA^ *vs*. Control^shRNA^ HeLa cell lysates (*n* = 3; Student’s *t*-test). G) Representative images of Alexa Fluoro 568-conjugated phalloidin-stained Control^shRNA^ and HD-PTP^shRNA^ HeLa cells transfected with EphB2-GFP, and stimulated 15’ with 1 μg/mL eB2 or Fc. H) Quantification of HeLa cell area shows that Control^shRNA^ EphB2-OE cells collapse to about half their size in response to eB2, while HD-PTP^shRN^ EphB2-OE cells collapse only by ~ 20% (*n* = 3, 60–80 cells/*n*; Control^shRNA^, *p* = 0.0003; HD-PTP^shRNA^, *p* = 0.0008; Student’s *t*-test). I) Representative images of Alexa Fluoro 568-conjugated phalloidin stains of Control^shRNA^ and HD-PTP^shRNA^ HeLa cells treated with 0.3 μg/mL Sema3A-Fc or Fc for 15’. J) Quantification of HeLa cell area shows Control^shRNA^ HeLa cells collapse to less than half their size in response to Sema3A-Fc, and HD-PTP^shRNA^ HeLa cells collapse to the same extent (*n* = 3, 60–80 cells/*n*; Control^shRNA^ *vs*. HD-PTP^shRNA^, *p* = 0.3880; Student’s *t*-test). Values are plotted as mean ± SD. All values can be found in Supplementary Table 4. kDa: kilodalton; eB2: ephrin-B2-Fc; *** *p* < 0.001; ** *p* < 0.01; * *p* < 0.05; n.s.: not significant. Scale bars: B) 15 μm, G and I) 10 μm. Inverted grayscale fluorescent images.

We next asked whether HD-PTP is involved in eB2-evoked cell collapse by transfecting Control^shRNA^ and HD-PTP^shRNA^ HeLa cells with an EphB2-GFP fusion expression plasmid (EphB2-OE) and stimulating them with eB2 or Fc. Similar transfection efficiency was confirmed in both cell types (Sup. Fig. 4A-C), but while Control^shRNA^ EphB2-OE cells treated with eB2 were decreased in size by ~50% compared to Fc-treated controls (Fig. 4G, H; *n* = 3; *p* = 0.0003), HD-PTP^shRNA^ EphB2-OE cell size was reduced by only 25% compared to controls (Fig. 4G, H; *n* = 3; Control^shRNA^ eB2 *vs*. HD-PTP^shRNA^ eB2, *p* = 0.0008). To determine whether this blunted response was specific to eB2 stimulation, Control^shRNA^ and HD-PTP^shRNA^ cells were exposed to Sema3A, another collapse-inducing chemotropic factor acting through neuropilin and plexin expressed by HeLa cells (Takahashi *et al.*, 1999; Thul *et al.*, 2017). Both cell lines collapsed to an equal extent when stimulated with Sema3A (Fig. 4I, J; *n* = 3; Control^shRNA^ *vs*. HD-PTP^shRNA^ *n.s., p* = 0.3880), indicating that the loss of HD-PTP blunts the eB2-induced collapse of HeLa cells, but does not affect the response to an unrelated cell collapse-inducing signal.

### HD-PTP expression and co-localisation with EphB2 in spinal motor neurons

Ephrin-B:EphB signalling is required for the guidance of embryonic spinal motor axons to their limb targets (Luria *et al.*, 2008; Poliak *et al.*, 2015), raising the possibility that HD-PTP may also be required for this process. First, we visualised *PTPN23* (gene encoding HD-PTP) mRNA in embryonic chick spinal cord at Hamburger and Hamilton stages (HH st.) 25 and 28, when spinal lateral motor column (LMC) axons are guided by ephrin-B:EphB signalling (Hamburger & Hamilton, 1951; Luria *et al.*, 2008). At these stages, *PTPN23* mRNA was expressed broadly in the dorsal spinal cord, including in motor columns identified by *ISL1* mRNA expression (Fig. 5A); however, mRNAs encoding the closely related phosphatases PTPN13 and PTPN14 were not detected (Fig. 5A). We also examined the relationship between EphB2 and HD-PTP expression levels and localisation in LMC neurons by electroporating EphB2-GFP and GFP-only expression plasmids into HH st. 18/19 chicken neural tubes (Kao *et al.*, 2009; Croteau & Kania, 2011) and explanting maturing LMC neurons at HH st. 25. There, we observed an approximately two-fold upregulation of HD-PTP protein in growth cones of EphB2-GFP-expressing neurons compared to GFP controls (Fig. 5E, G; *n* = 3; *p* = 0.0077) and preferential co-localisation of HD-PTP with EphB2-GFP (Fig. 5H, I; *n* = 3; *p* = 0.0153).

**Figure 5.**
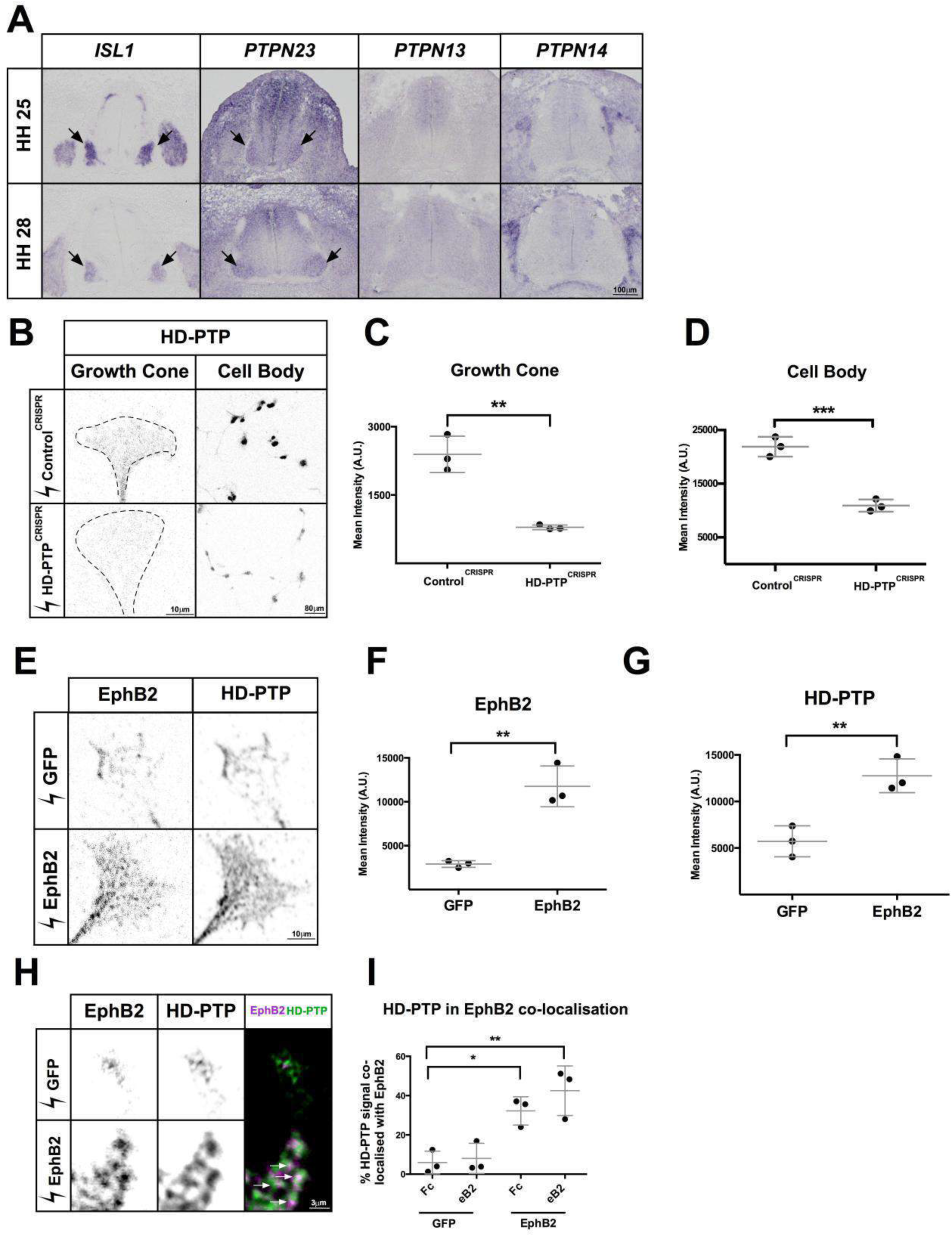
HD-PTP expression and co-localisation with EphB2 in embryonic motor neurons. A) Representative images of chick embryonic spinal cord sections at HH st. 25 and HH st. 28 where *ISL1, PTPN23* (chicken HD-PTP-encoding gene), *PTPN13* and *PTPN14* mRNA was detected using *in situ* hybridisation. Note expression of *PTPN23* in *ISL1*-expressing motor column (arrows). B) Representative images of anti-HD-PTP antibody staining in growth cones and cell bodies of dissociated motor neurons harvested from embryonic spinal cords electroporated with Control^CRISPR^ or HD-PTP^CRISPR^ plasmids. C) Quantification of HD-PTP signals in growth cones of dissociated motor neurons harvested from embryonic spinal cords shows a decreased signal in HD-PTP^CRISPR^ compared to Control ^CRISPR^ (*n* = 3, 10–12 growth cones/*n*; *p* = 0.0023; Student’s *t*-test). D) Quantification of HD-PTP signal in cell bodies of dissociated motor neurons harvested from embryonic spinal cords show decrease signal in HD-PTP^CRISPR^ compared to Control^CRISPR^ (*n* = 3, 30–50 cell bodies/*n*; *p* = 0.0009; Student’s *t*-test). E) Representative images of anti-EphB2 and anti-HD-PTP antibody staining of growth cones of dissociated motor neurons harvested from embryonic spinal cords electroporated with GFP- or EphB2-GFP-expressing plasmids. Up-regulation of HD-PTP expression is evident in EphB2 over-expressing growth cones. F) Quantification of EphB2 signal in growth cones of dissociated motor neurons harvested from embryonic spinal cords show an increased signal in EphB2-GFP compared to GFP (*n* = 3, 10–12 growth cones/*n*; *p* = 0.0077; Student’s *t*-test). G) Quantification of HD-PTP signal in growth cones of dissociated motor neurons harvested from embryonic spinal cords show an increased signal in EphB2-GFP compared to GFP (*n* = 3, 10–12 growth cones/*n*; Student’s *t*-test). H) Representative images of growth cones of dissociated motor neurons harvested from embryonic spinal cords electroporated with GFP- or EphB2-GFP-expressing plasmids showing co-localisation of HD-PTP and EphB2 (arrows). I) Quantification of HD-PTP signal localisation in EphB2-positive puncta in growth cones of dissociated motor neurons harvested from embryonic spinal cords electroporated with GFP- or EphB2-GFP-expressing plasmids. HD-PTP is preferentially found in EphB2-containing sites in EphB2-GFP growth cones compared to GFP-expressing growth cones (*n* = 3, 10–12 growth cones/*n*; *p* = 0.0153; Student’s *t*-test). Values are plotted as mean ± SD. All values can be found in Supplementary Table 4. eB2: ephrin-B2-Fc; *** *p* < 0.001; ** *p* < 0.01; * *p* < 0.05. Scale bars: A) 100 μm, B) 10 μm and 80 μm, E) 10 μm, H) 3 μm. Inverted grayscale fluorescent images except for visible light images in A and dual colour images in H.

### HD-PTP is required for ephrin-B2-induced LMC growth cone collapse

To test whether HD-PTP is required for normal ephrin-B:EphB signalling in LMC neurons, we induced HD-PTP loss-of-function in LMC motor neurons using CRISPR-Cas9 (Cong *et al.*, 2013; Shinmyo *et al.*, 2016). We designed three guide RNAs targeting exons 2 to 5 of the *PTPN23* gene to increase the likelihood of coding sequence double-stranded breaks and frameshifts due to Cas9 error-prone non-homologous end joining (Sup. Fig. 5A; Véron *et al.*, 2015; Doudna & Charpentier, 2014). We co-electroporated three plasmids, each encoding one guide RNA, a Cas9-FLAG fusion protein, and GFP expressed using the T2A self-cleaving peptide system, into HH st. 18/19 chick neural tubes and harvested HD-PTP^CRISPR^ spinal cords at HH st. 25. As a control, we used a plasmid encoding Cas9-GFP-FLAG and a guide RNA targeting an untranslated region of the *EPHA4* gene (Control^CRISPR^). PCR amplification of genomic DNA extracted from HD-PTP^CRISPR^, but not from Control^CRISPR^ spinal cords revealed the presence of a deletion in the *PTPN23* locus consistent with a deletion between guides 1 and 3 (Sup. Fig. 5B). HD-PTP signal in cultured HD-PTP^CRISPR^ LMC growth cones and cell bodies was significantly decreased compared to Control^CRISPR^ controls (Fig. 5B-D; *n* = 3; growth cone, *p* = 0.0023; cell body, *p* = 0.0009).

Explanted HH st. 25 HD-PTP^CRISPR^ and Control^CRISPR^ LMC neurons, dissociated and cultured for at least 18 hours, did not differ in their capacities to form growth cones (Fig. 5B), extend axons (Sup. Fig. 6A), or express EphB2 (Sup. Figs. 5E, F). To determine whether HD-PTP is required for LMC growth cone eB2:EphB2 signalling, we focused on the medial subpopulation of LMC neurons, which express high levels of EphB2 and are repelled by eB2 *in vivo* and *in vitro* (Luria *et al.*, 2008; Kao & Kania, 2011). HD-PTP^CRISPR^ or Control^CRISPR^ LMC neurons were dissociated and medial LMC neurons were identified by the expression of the transcription factor Isl1 (Tsuchida *et al.*, 1994). Control^CRISPR^ medial LMC growth cones collapsed significantly when treated with eB2, but HD-PTP^CRISPR^ medial LMC growth cones showed a markedly attenuated collapse response (Fig. 6A, D; *p* < 0.0001). This effect was specific to eB2 treatment, since HD-PTP^CRISPR^ and Control^CRISPR^ growth cones collapsed to the same extent when exposed to Sema3F, a protein known to repel medial LMC axons (Fig. 6B, D; Huber *et al.*, 2005).

**Figure 6.**
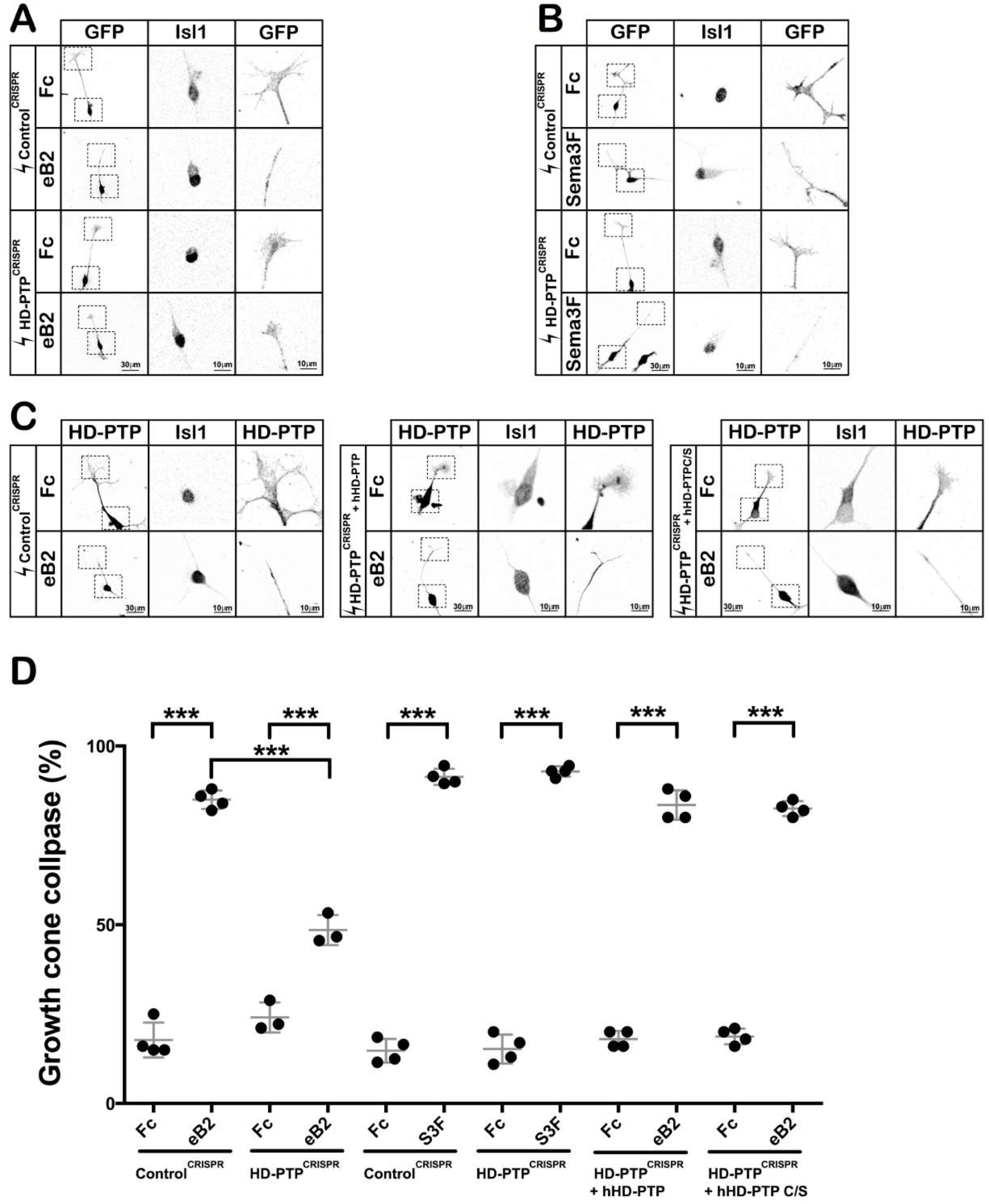
Spinal motor axon growth cones require HD-PTP for ephrin-B2-induced collapse. A) Representative images of GFP^+^ neurons from dissociated Control^CRISPR^- or HD-PTP^CRISPR^-electroporated motor neurons, incubated with eB2 or Fc and stained with anti-GFP and anti-Isl1 antibodies. Insets show medial LMC Isl1-expressing cell bodies and growth cones. B) Representative images of GFP^+^ neurons from dissociated Control^CRISPR^- or HD-PTP^CRISPR^-electroporated motor neurons, incubated with Sema3F or Fc and stained with anti-GFP and anti-Isl1 antibodies. Insets show medial LMC Isl1-expressing cell bodies and growth cones. C) Representative images of rescue experiments with dissociated motor neurons electroporated with Control^CRISPR^ plasmid or HD-PTP^CRISPR^ co-electroporated with hHD-PTP or hHD-PTP C/S plasmid, incubated 30’ with 10 μg/mL eB2 or Fc and stained with anti-HD-PTP and anti-Isl1 antibodies. Insets show medial LMC Isl1-expressing cell bodies and growth cones. D) Quantification of collapsed growth cones in dissociated motor neurons electroporated with CRISPR constructs and stimulated with ligands. HD-PTP^CRISPR^ or Control^CRISPR^ growth cones were incubated for 30’ with 10 μg/mL eB2 or Fc. The collapse response of HD-PTP^CRISPR^ growth cones to eB2 was significantly attenuated compared to Control^CRISPR^ (*n* = 3, 90 growth cones/*n*; *p* < 0.0001; Fisher’s exact test). HD-PTP^CRISPR^ or Control^CRISPR^ growth cones were incubated for 30’ with 0.3 μg/mL Sema3F-Fc or Fc. The two CRISPR growth cone populations behaved identically, demonstrating that HD-PTP loss does not affect the response to Sema3F (*n* = 4, 30 growth cones/*n*; Fisher’s exact test). Rescue experiments with growth cones from dissociated motor neurons electroporated with Control^CRISPR^, or HD-PTP^CRISPR^ co-electroporated with hHD-PTP or hHD-PTP C/S expression plasmid, incubated 30’ with 10 μg/mL eB2 or Fc and stained with anti-HD-PTP and anti-Isl1 antibodies. Both populations responded to eB2 treatment indistinguishably from control (*n* = 4, 50 growth cones/*n*; Fisher’s exact test). Values are plotted as mean ± SD. All values can be found in Supplementary Table 4. h: human; S3F: Sema3F; eB2: ephrin-B2-Fc; *** *p* < 0.001; n.s.: not significant. Scale bars: A-C) 30 μm, insets 10 μm. Inverted grayscale fluorescent images.

To further characterise the specificity of the HD-PTP knockdown, we carried out rescue experiments by co-electroporating a human (h) HD-PTP expression plasmid together with the chick-specific HD-PTP^CRISPR^ plasmids as above. In medial LMC neurons co-electroporated with HD-PTP^CRISPR^ and hHD-PTP expression plasmids, HD-PTP protein levels returned close to control levels (Sup. Fig. 6B, C), as did their collapse response to eB2 (Fig. 6C, D). We also asked whether the HD-PTP phosphatase domain is required for its function in growth cone collapse by co-electroporating HD-PTP^CRISPR^ plasmids together with a plasmid encoding a human HD-PTP with a phosphatase active site-disrupting point mutation (hHD-PTP C/S; Cao *et al.*, 1998). This mutant HD-PTP was capable of rescuing the HD-PTP^CRISPR^–induced growth cone collapse defect (Fig. 6C, D), suggesting that HD-PTP requirement for eB2 growth cone collapse is very likely phosphatase activity-independent.

### HD-PTP is required for ephrin-B2:EphB2-mediated medial LMC guidance in vivo

We next hypothesised that HD-PTP loss in LMC neurons *in vivo* would lead to an abnormal medial LMC axons entry into the dorsal limb nerve, similar to a phenotype observed in mice with a genetic loss of ephrin-B:EphB signalling (Luria *et al.*, 2008). To visualise the axons of medial LMC neurons, we co-electroporated them with the HD-PTP^CRISPR^ or Control^CRISPR^ guide expression plasmids lacking the GFP, and the medial LMC-specific axonal marker plasmid *e[Isl1]∷GFP* (Kao *et al.*, 2009; Fig. 7D). Loss of HD-PTP function did not result in abnormal LMC neuron specification or survival at HH st. 25, when LMC axons enter the dorsal and ventral hindlimb nerves (Fig. 7A-C; Landmesser, 2018). At this stage, in Control^CRISPR^ + *e[Isl1]GFP* embryos, 7% of axonal GFP signal was found in the dorsal nerve and 93% in the ventral nerve, similar to the incidence of medial LMC labelling by retrograde fill from dorsal and ventral limb muscles (Luria *et al.*, 2008). In contrast, in HD-PTP^CRISPR^ + *e[Isl1]∷GFP* embryos, ~25% of axonal GFP signals were found in the dorsal nerve and ~75% of them were found in the ventral nerve, a significant difference from controls (Fig. 7E, F; *n* = 5; *p* = 0.0149), demonstrating that HD-PTP is required for the normal guidance of medial LMC motor axons *in vivo*.

**Figure 7.**
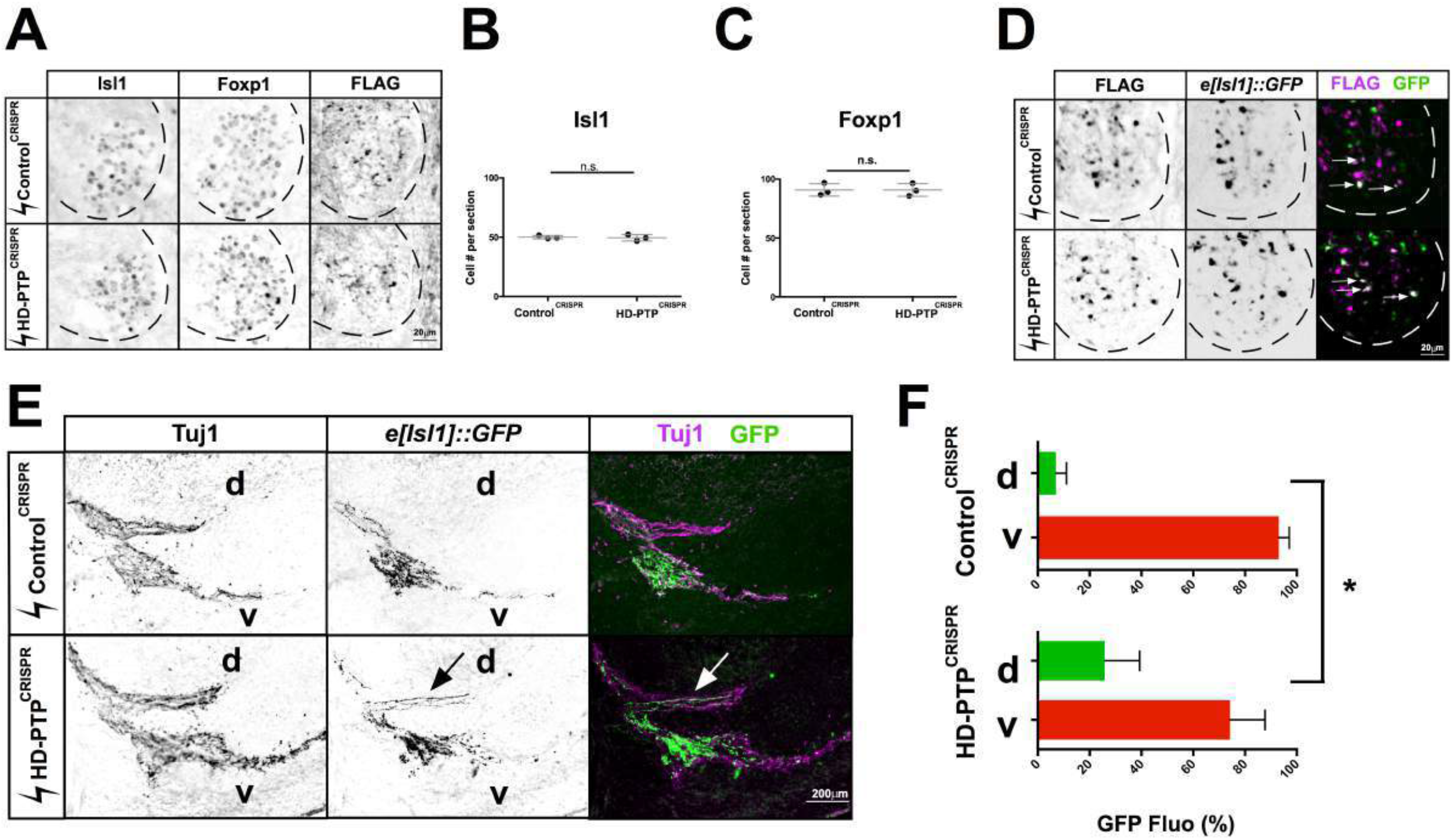
HD-PTP is required for ephrin-B:EphB-mediated motor axon guidance *in vivo*. A) Representative sections Control^CRISPR^ and HD-PTP^CRISPR^ HH St. 25 spinal cords showing expression of Isl1, Foxp1 and FLAG, the Cas9 expression marker, demonstrating efficient electroporation of motor neurons. B) Quantification of Isl1+ medial LMC neurons in Control^CRISPR^ and HD-PTP^CRISPR^ embryos. Their numbers are not significantly different between the two populations of embryos (*n* = 3, 10 sections/*n*; Student’s *t*-tests). C) Quantification of Foxp1+ LMC motor neurons in Control^CRISPR^ and HD-PTP^CRISPR^ embryos. Their numbers do not differ between the two conditions (*n* = 3, 10 sections/*n*; Student’s *t*-tests). D) Representative images of the FLAG Cas9 expression marker and the medial LMC marker in *e[Isl1]∷GFP* in Control^CRISPR^ and HD-PTP^CRISPR^ sections of HH St. 25 ventral spinal cords. E) Representative images of the limb nerve in Control^CRISPR^ and HD-PTP^CRISPR^ HH St. 25 embryos, stained with anti-Tuj1 antibodies to reveal limb nerves and *e[Isl1]∷GFP* in medial LMC axons. Medial axons aberrantly innervate the dorsal mesenchyme in HD-PTP^CRISPR^ embryos. F) Quantification of *e[Isl1]∷GFP* expression in dorsal vs. ventral nerves. Control^CRISPR^ embryos contain ~93% of GFP in the ventral nerve and ~7% in the dorsal nerve. HD-PTP^CRISPR^ embryos contain ~74% of GFP in the ventral nerve and ~26% in the dorsal nerve, demonstrating that disruption of HD-PTP *in vivo* disrupts the fidelity of misrouted medial LMC axon projection (*n* = 5 embryos, 10–20 sections/*n*; *p* = 0.0149; Student’s *t-* tests between GFP signal % in dorsal Control^CRISPR^ vs. dorsal HD-PTP^CRISPR^). Values are plotted as mean ± SD. All values can be found in Supplementary Table 4. d: dorsal; v: ventral; * *p* < 0.05; n.s.: not significant. Scale bars: B-C) 20 μm, D) 200 μm. Inverted grayscale fluorescent images except for dual colour images in E.

### HD-PTP is required for ephrin-B2-induced EphB2 phosphorylation, SFK activation, and EphB2 surface patching

Eph forward signalling is a multi-step process, involving the phosphorylation of Src Family Kinases (SFKs) on their activating tyrosine, Y418 (Knöll & Drescher, 2004; Poliak *et al.*, 2015). To examine whether this step requires HD-PTP, we used an antibody specific for this phosphorylation (Boggon & Eck, 2004), in HeLa cells and medial LMC growth cones with decreased HD-PTP expression, exposed to eB2 or Fc (Fig. 8C-F). Control^shRNA^ EphB2-OE HeLa cells stimulated with eB2 had an almost three-fold increase in phospho-Y418-SFK signal compared to Fc stimulation (Fig. 8C top, D; *p* = 0.0227). However, HD-PTP^shRNA^ EphB2-OE HeLa cells had no detectable change in phospho-Y418-SFK signal upon ligand treatment when compared to Fc (Fig. 8C bottom, D; *p* = 0.7109). Similarly, Control^CRISPR^ LMC growth cones treated with eB2 displayed increased levels of phospho-Y418-SFK signal compared to Fc-treated growth cones (Fig. 8E top, F; *n* = 3; *p* < 0.0001; Poliak *et al.*, 2015), while HD-PTP^CRISPR^ LMC growth cones did not show this effect (Fig. 8E bottom, F; *p* = 0.9810). Thus, the loss of HD-PTP abolishes ephrin-B2-induced activation of a critical EphB2 effector.

**Figure 8.**
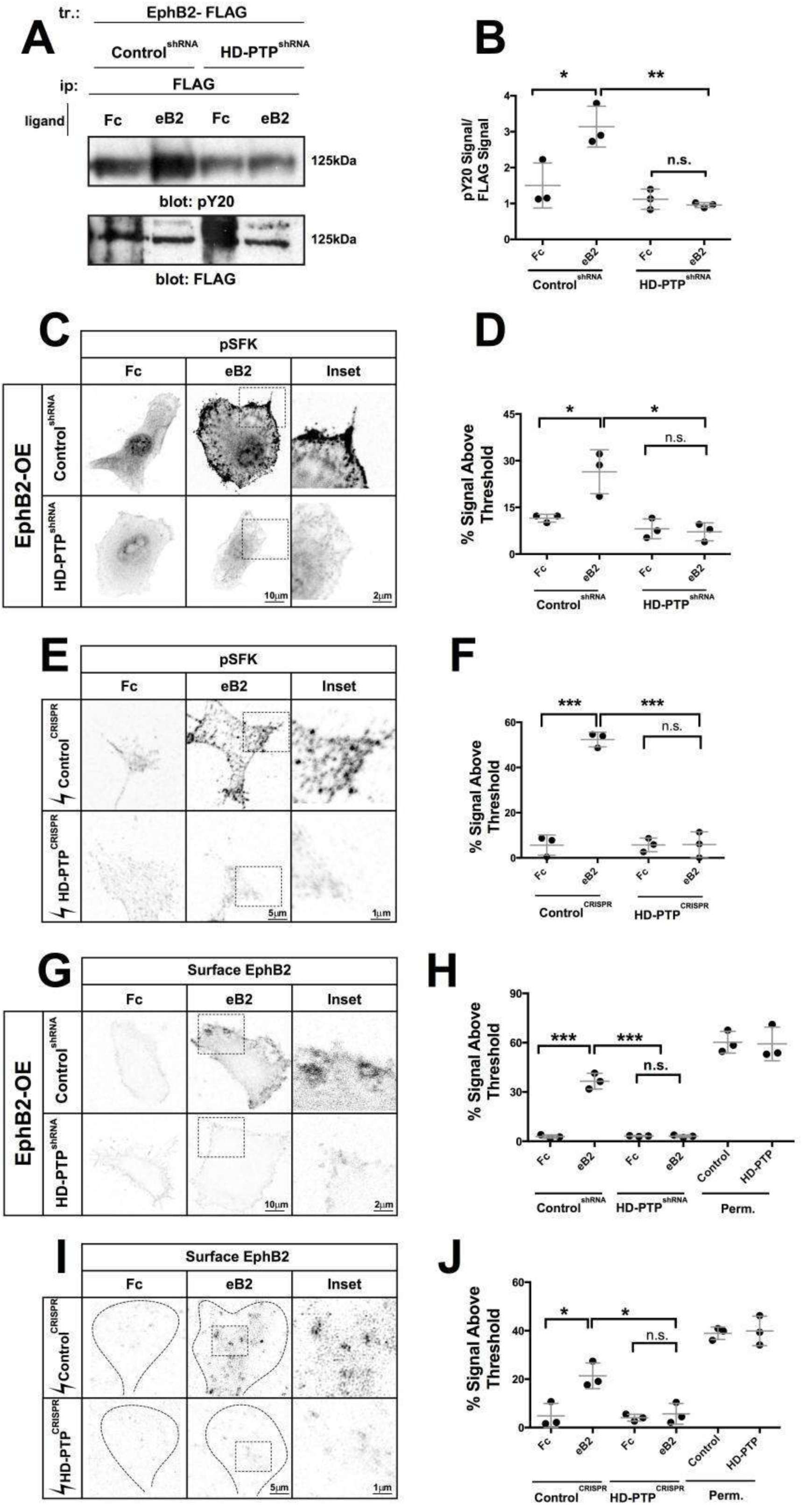
HD-PTP is required for ephrin-B-induced EphB2 phosphorylation, SFK phosphorylation, and EphB2 surface patching. A) Representative Western blot using anti-phosphotyrosine and anti-FLAG antibodies after pull-downs of EphB2 (with anti-FLAG antibodies) in Control^shRNA^ and HD-PTP^shRNA^ HeLa cells, stimulated with 1 μg/mL eB2 or Fc for 5’. The band size corresponds to EphB2. B) Quantification of phosphotyrosine signal over FLAG signal shows ligand-induced phosphorylation of EphB2 in Control^shRNA^ HeLa cells (*p* = 0.0284), but not in HD-PTP^shRNA^ cells (*p* = 0.3908) (*n* = 3; one-way ANOVA followed by Student’s *t*-tests). C) Representative images of Control^shRNA^ and HD-PTP^shRNA^ HeLa cells, incubated for 5’ with 1 μg/mL eB2 or Fc and stained with anti-phospho-Y418-SFK antibodies showing increased SFK activation following eB2 exposure. D) Quantification of anti-phospho-Y418-SFK staining in Control^shRNA^ and HD-PTP^shRNA^ HeLa cells incubated for 5’ with 1 μg/mL eB2 or Fc. Control^shRNA^ showed an increase in phopho-Y418-SFK signal upon eB2 stimulation (*p* = 0.0227), yet HD-PTP^shRNA^ HeLa cells display no detectable increase in SFK phosphorylation (*p* = 0.7109) (*n* = 3, 10–12 cells/*n*; one-way ANOVA followed by corrected Student’s *t*-tests). E) Representative images of Control^CRISPR^ and HD-PTP^CRISPR^ spinal motor neuron growth cones, incubated for 15’ with 10 μg/mL eB2 or Fc and stained for with anti-phospho-Y418-SFK, revealing SFK activation following eB2 exposure. F) Quantification of anti-phospho-Y418-SFK signal in Control^CRISPR^ and HD-PTP^CRISPR^ motor neuron growth cones, incubated for 15’ with 10 μg/mL eB2 or Fc. Control^CRISPR^ growth cones showed a ligand-induced increase in SFK activation (*p* < 0.0001), but HD-PTP^CRISPR^ growth cones did not (*p* = 0.9810) (*n* = 3, 10–12 growth cones/*n*; one-way ANOVA followed by corrected Student’s *t*-tests). G) Representative images of Control^shRNA^ and HD-PTP^shRNA^ shRNA HeLa cells, incubated for 5’ with 1 μg/mL eB2 or Fc and immunostained for EphB2 using a non-permeabilising fixation conditions (see methods and Supplemental Fig. 8). EphB2 patching is visualised through increased signal intensity of surface EphB2 staining. H) Quantification of surface EphB2 patching in Control^shRNA^ and HD-PTP^shRNA^ HeLa cells, incubated for 5’ with 1 μg/mL eB2 or Fc, measured by percentage of the cell area containing anti-EphB2 signal. In stark contrast to Control^shRNA^ cells (*p* = 0.0003), HD-PTP^shRNA^ HeLa cells failed to elicit EphB2 surface patching upon ligand binding (*p* = 0.8609) (*n* = 3, 10–12 cells/*n*; one-way ANOVA followed by corrected Student’s *t*-tests). I) Representative images of Control^CRISPR^ and HD-PTP^CRISPR^ motor neuron growth cones, incubated for 15’ with 10 μg/mL eB2 or Fc and immunostained for EphB2 using non-permeabilising fixation conditions. EphB2 patching is visualised through increased signal intensity of surface anti-EphB2 staining. J) Quantification of EphB2 patching in Control^CRISPR^ and HD-PTP^CRISPR^ motor neuron growth cones, incubated for 15’ with 10 μg/mL eB2 or Fc, as measured by percentage of the growth cone area containing surface EphB2 signal. In contrast to Control^CRISPR^ growth cones (*p* = 0.017), HD-PTP^CRISPR^ growth cones failed to elicit EphB2 surface patching upon ligand binding (*p* = 0.5707) (*n* = 3; one-way ANOVA followed by corrected Student’s *t*-tests). Values are plotted as mean ± SD. All values can be found in Supplementary Table 4. kDa: kilodalton; eB2: ephrin-B2-Fc; *** *p* < 0.001; * *p* < 0.05; n.s.: not significant. Scale bars: C and G) 10 μm, inset 2 μm, E, I) 5 μm, inset 1 μm. Inverted grayscale fluorescent images except for dual colour images in J.

Another early step in the Eph signalling cascade is the phosphorylation of a juxtamembrane tyrosine residue critical for Eph kinase activity (Zisch *et al.*, 1998). To determine whether HD-PTP is important for this, we transfected Control^shRNA^ and HD-PTP ^shRNA^ HeLa cells with an EphB2-FLAG expression plasmid and stimulated with eB2 or Fc. We then lysed the cells and performed anti-FLAG pull-down and immunoblotted with an anti-phosphotyrosine antibody. Compared to Fc treatment, a significant increase in EphB2 phosphorylation was observed following eB2 stimulation in Control^shRNA^ cells (Fig. 8A, B; *n* = 3; *p* = 0.0284); however, this effect was absent in HD-PTP ^shRNA^ cells (Fig. 8A, B; *n* = 3; *p* = 0.3908).

One of the first events of ephrin-Eph signalling is the formation of receptor-ligand multimer arrays on the cell surface (Torres *et al.*, 1998; Seiradake *et al.*, 2013; Schaupp *et al.*, 2014), raising the possibility that this process is HD-PTP dependent. Immunohistochemical detection of cell surface EphB2 in Control^shRNA^ EphB2-OE HeLa cells, revealed that eB2 treatment resulted in significant EphB2 cell surface patch formation, a correlate of Eph receptor multimers (Fig. 8G top, H; *p* = 0.0003 *v*. Fc; Seiradake *et al.*, 2013). In contrast, HD-PTP^shRNA^ EphB2-OE HeLa cells showed a conspicuous absence of eB2-induced EphB2 cell surface patches compared to Fc treatment (Fig. 8G bottom, H; *p* = 0.8609). Similarly, Control^CRISPR^ LMC growth cones treated with eB2 showed increased EphB2 surface signal patching compared to Fc-treated ones (Fig. 8I top, J; *p* = 0.017). In contrast, HD-PTP^CRISPR^ LMC growth cones did not display such effects (Fig. 8I bottom, J; *p* = 0.5707), suggesting a critical role for HD-PTP in eB2-induced surface clustering of EphB2.

### HD-PTP protects EphB2 from ligand-induced degradation

As a component of the ESCRT complex, HD-PTP controls the endocytic pathway degradation of ligand-bound cell surface receptors as well as their salvage through recycling endosomes (Ichioka *et al.*, 2007; Doyotte *et al.*, 2008). To examine whether HD-PTP may direct such processing of Eph receptors, we compared EphB2 protein levels following protein synthesis inhibition in the presence or absence of eB2, in cells with diminished HD-PTP levels. To do this, we transfected EphB2-FLAG expression plasmids into Control^shRNA^ and HD-PTP^shRNA^ HeLa cells, treated them with eB2 or Fc in the presence of the protein synthesis inhibitor cycloheximide (CHX) and measured dynamic changes in EphB2 protein levels via FLAG immunoblotting (Kharitidi *et al.*, 2015). Fc-treated Control^shRNA^ HeLa cells maintained a steady level of EphB2 until about 30 minutes after CHX addition, when EphB2 levels began to decrease (Fig. 9A, B). When incubated with eB2 and CHX, however, EphB2 levels remained steady for up to 60 minutes (Fig. 9A, B, J; *p* = 0.0018), suggesting that eB2 exposure may inhibit EphB2 degradation. In contrast, 30 minutes after Fc and CHX exposure, HD-PTP^shRNA^ HeLa cells had lower EphB2 levels compared to Fc-treated Control^shRNA^ HeLa cells (Fig. 9A-D; *n* = 3; *p* = 0.002). Furthermore, eB2 and CHX treatment of HD-PTP^shRNA^ HeLa cells resulted in an even more rapid decrease of EphB2 levels, with their almost complete depletion after 60 minutes of treatment (Fig. 9C, D; *n* = 3; 60-min Control^shRNA^ eB2 vs. 60-min HD-PTP^shRNA^ eB2, *p* = 0.0038). Together, these experiments suggest that, in contrast to HD-PTP loss leading to an increase in levels of other tyrosine kinase receptors, HD-PTP silencing accelerates EphB2 depletion following ligand exposure.

**Figure 9.**
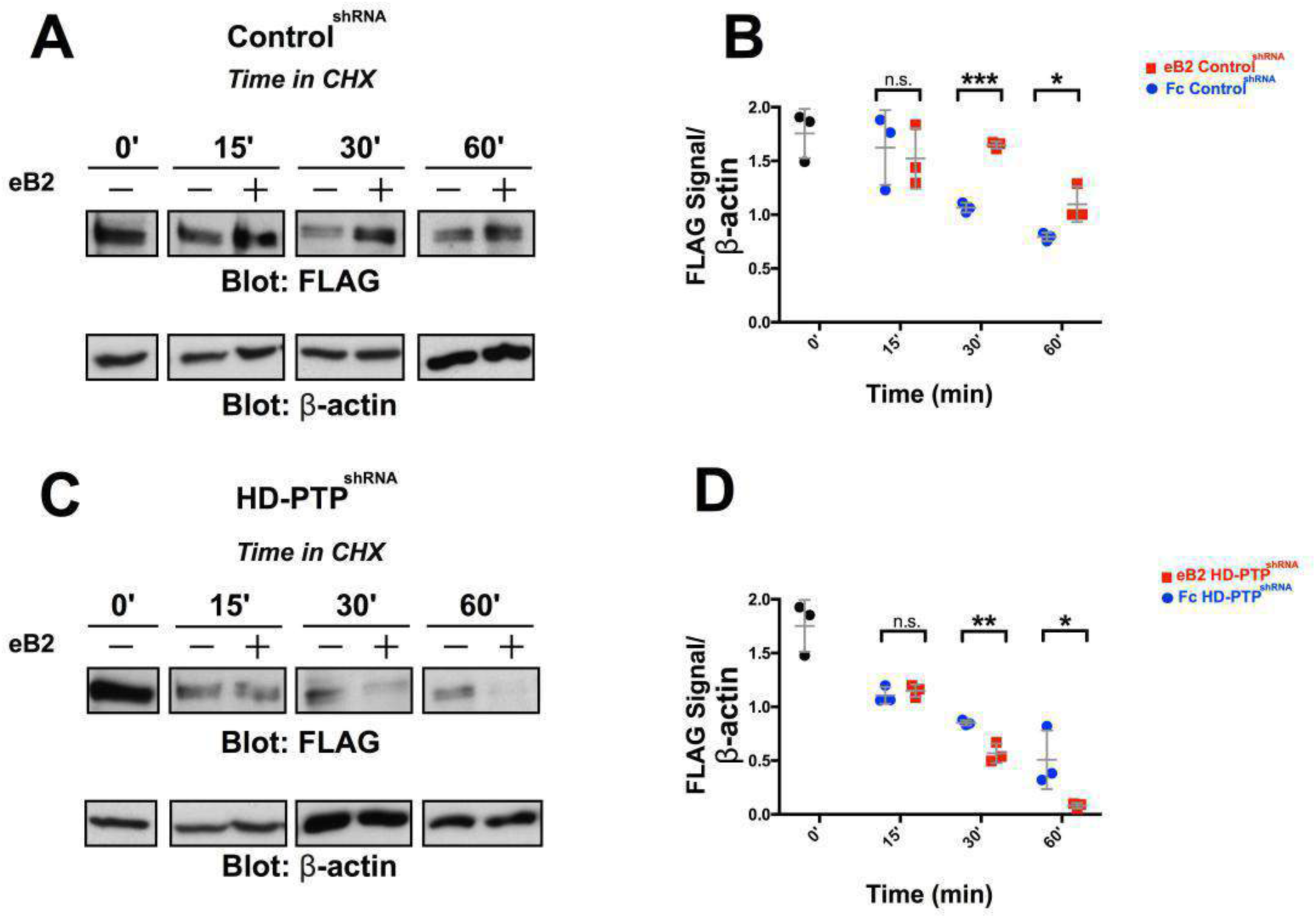
HD-PTP loss increases the rate of EphB2 degradation. A) Representative Western blot for EphB2 expression detected with anti-FLAG antibodies in transfected Control^shRNA^ HeLa cell lysates at different time points after incubation with 10 μg/mL protein synthesis blocker cycloheximide, exposed to either 1 μg/mL eB2 or Fc. β-actin detection is used as an internal control. B) Quantification of Western blots for EphB2 detected with anti-FLAG antibodies in transfected Control^shRNA^ HeLa cell lysates after incubation with 10 μg/mL protein synthesis blocker cycloheximide together with either 1 μg/mL eB2 or Fc. FLAG signal intensity was normalised to β-actin and plotted for the different time points. By 30’ after cycloheximide treatment, eB2 stimulation appears to protect EphB2 from degradation compared to Fc (*n* = 3; Student’s *t*-test). C) Representative Western blot for EphB2 detected with anti-FLAG antibodies in transfected HD-PTP^shRNA^ HeLa cell lysates at different time points after incubation with 10 μg/mL protein synthesis blocker cycloheximide and either 1 μg/mL eB2 or Fc. β-actin is used as an internal control. D) Quantification of Western blots for EphB2 detected with anti-FLAG antibodies in transfected HD-PTP^shRNA^ HeLa cell lysates after incubation with 10 μg/mL protein synthesis blocker cycloheximide together with either 1 μg/mL eB2 or Fc. FLAG signal intensity was normalised to β-actin and plotted for the different time points. In contrast to Control^shRNA^ HeLa cells, in HD-PTP^shRNA^ HeLa cells, eB2 stimulation appears to increase rate of EphB2 degradation compared to Fc by 30’ after cycloheximide treatment (*n* = 3; Student’s *t*-test). Values are plotted as mean ± SD. All values can be found in Supplementary Table 4. CHX: cycloheximide; eB2: ephrin-B2-Fc; *** *p* < 0.001; ** *p* < 0.01; * *p* < 0.05; n.s.: not significant.

## Discussion

Our proteomics experiments identify a number of potential novel effectors of Eph signalling, and demonstrate that one such protein is the ESCRT adaptor HD-PTP. Its association with EphB2 is potentiated by ephrin-B2 binding, and its function is required for repulsive responses to ephrin-B2 in cultured cells and motor neuron growth cones, as well as the normal guidance of spinal motor axons *in vivo*, a process that relies on ephrin-B:EphB signalling. In addition to being essential for the earliest step of ephrin-B:EphB signalling, HD-PTP also protects EphB2 against ligand-induced degradation. Here, we discuss these findings in the context of general principles of ephrin:Eph signalling and the role of ESCRT proteins in this process, as well as in axon guidance and other Eph functions.

### Insights into Eph signalling revealed by BioID

We used BioID and mass spectrometry to describe the EphB2-associated protein landscape during forward signalling. Our list of EphB2-proximal proteins includes some known EphB2 effectors such as NCK1, NCK2, CRK, and YES, arguing that our ephrin-B2 ligand differential strategy identifies biologically-relevant protein-protein interactions (Fawcett *et al.*, 2007; Hock *et al.*, 1998; Zisch *et al.*, 1998). Given that ephrin-evoked Eph signalling is a short-lived event, occurring on the scale of minutes, and that our biotinylation of EphB2-proximal proteins proceeded on the scale of hours, our results suggest that the BioID methodology used in these experiments is able to capture even relatively short-lived protein-protein interactions. Beyond specific protein hits, our biological process and pathway analysis results align with previously defined functions of ephrin:Eph signalling in neurodevelopment and cytoskeletal organisation, through its action at cell-cell junctions, cell periphery and membrane, and GTPase regulation (Kania & Klein, 2016). In the context of EphB-mediated axon guidance, our data confirm the association of EphB2 with the Unc5 class of netrin receptors, which results in synergistic EphB signalling (Poliak *et al.*, 2015). Furthermore, together with the BioID data set of EphA2-proximal proteins (White *et al.*, 2017), our results point to several biological processes in ephrin:Eph signalling that lack a detailed mechanistic description: endosomal transport and cell division and differentiation. Endocytosis plays a prominent role in Eph signalling (Pitulescu & Adams, 2010), and ESCRT components have been previously associated with Eph receptors, although only in the context of Eph-containing exosomes (Gong *et al.*, 2016). In addition to HD-PTP, another endosome-associated protein apparently recruited to activated EphB2 is the RUN and FYVE domain-containing protein 1 (RUFY1), whose knockdown phenotype suggests a function with HD-PTP in EGFR trafficking (Gosney *et al.*, 2018). While ephrin:Eph signalling has been previously linked to cell differentiation and proliferation, the mechanism of this is not well understood because few direct ephrin:Eph effectors of these functions have been identified (Genander & Frisén, 2010). This aspect of Eph signalling has been explored at the transcriptome level, implicating the PI3-Kinase and Abl-cyclinD1 pathways, and more recently, histone methylation via Akt (Genander *et al.*, 2009; Fawal *et al.*, 2018). Our proteomic identification of EphB2 proximal proteins as Abl2 and Pik3r1, confirms these links, but also suggests that Notch2 may be a novel intermediary that allows Eph receptors to intersect with a transcriptional response pathway controlling a multitude of developmental and homeostatic processes (Andersson *et al.*, 2011).

### HD-PTP: a new and potent effector of Eph signalling

Our experiments show that HD-PTP can form an ephrin-B2-driven complex with EphB2 and plays a critical and early role in Eph signalling: cells and growth cones with even a partial loss of HD-PTP exhibit a marked disruption of collapse responses to ephrin-B, apparently because of decreased Eph receptor clustering, phosphorylation and activation of Src family kinases. Among these, Eph receptor clustering is the most upstream event following ephrin-B2 binding, and a defect at either of these steps could explain reduced downstream phosphorylation. Since HD-PTP gain- or loss-of-function does not affect EphB2 abundance or surface localisation, the simplest explanation of these effects would place HD-PTP at the ephrin-B2:EphB2 binding and/or clustering steps of signalling. These steps depend on the extracellularly-located ligand-binding and the cysteine-rich domains (Smith *et al.*, 2004; Himanen *et al.*, 2010; Seiradake *et al.*, 2010), but are also modulated by the intracellular PDZ and SAM domains whose deletion enhances ephrin-induced Eph clustering and signalling in cultured cells (Schaupp *et al.*, 2014). Although without extensive biochemical analysis we are unable to resolve between a role of HD-PTP in ligand binding or receptor clustering, this study is the first report identifying an intracellular protein whose loss has a profound impact on the earliest steps of Eph receptor activation. Of course, one possible explanation for such strong phenotypes could be that, in line with HD-PTP’s role in ESCRT pathway progression, HD-PTP’s loss affects indirectly the expression or subcellular localisation of a protein required for the early steps of Eph signalling. Given that semaphorin-mediated cellular responses are normal under HD-PTP loss-of-function conditions, such an indirect effect would have to be specific to the Eph signalling pathway. Nevertheless, since no intracellular proteins essential for binding or clustering of Eph receptors have been identified, and considering the impact of HD-PTP loss on the earliest molecular events of Eph signalling, HD-PTP either a regulates a potent factor required for the binding and/or clustering steps, or is participating in these steps directly. Because of its ability to interact with EphB2, we favour the latter possibility; either way, our data argue that HD-PTP is an important molecular handle on the mechanisms regulating of the earliest steps of Eph signalling.

Upon ligand exposure, cells with a loss of HD-PTP display increased degradation of EphB2 receptor, a surprising observation given HD-PTP’s known function in promoting the progression of activated cell surface receptors through the endocytic pathway. For example, it has been shown to attenuate intracellular signalling downstream of ligand-activated cell surface receptors such as integrin α5β1, E-cadherin, and EGFR, such that a loss of HDP-PTP results in the endocytic accumulation of these receptors and their exacerbated signalling (Kharitidi *et al.*, 2015; Lin *et al.*, 2011; Doyotte *et al.*, 2008). Our results suggest that in the context of Eph receptor function, HD-PTP promotes signalling. How may it do that? Some insights come from the observation of reduced Wnt signalling due to the impaired function of the Wnt receptor in *Drosophila* wing imaginal disks lacking HD-PTP (Pradhan-Sundd & Verheyen, 2015). This study argues that HD-PTP recruits deubiquitylases to counterbalance the ubiquitylation of both Wnt receptor and the endosome-associated protein Hrs/HGS, which normally promotes the recycling of Wnt receptors destined for lysosomal degradation. In this context, HD-PTP loss results in increased ubiquitylation and lysosomal degradation of Hrs/HGS, leading to increased endocytic accumulation of ubiquitylated Wnt receptors and their decreased recycling. Thus, one explanation for the accelerated depletion of EphB2 following ephrin-B stimulation observed in HD-PTP deficient cells could be through a similar impact on deubiquitylation of endosomal proteins or Eph receptors themselves. Indeed, two studies show that EphB receptors are ubiquitylated in response to ligand binding. EphB2 is ubiquitylated by the SOCS box protein SPSB4, whose loss accentuates ephrin-B2-induced repulsive cellular responses (Okumura *et al.*, 2017), and the ligand-induced kinase activity of EphB1 promotes its ubiquitylation by Cbl (Fasen *et al.*, 2008). Therefore, an alternative model of HD-PTP function may involve its recruitment of deubiquitylases that counterbalance ephrin-B-induced EphB ubiquitylation, as well as promoting its recycling, since deubiquitylated Eph receptors are more likely to be recycled (Sabet *et al.*, 2015). Together, our data suggest that HD-PTP has dual roles in the ephrin-B:EphB signalling cascade: early on, it is required for the initiation of signalling, and further downstream, it acts as a negative regulator of receptor degradation (Fig. 10).

**Figure 10.**
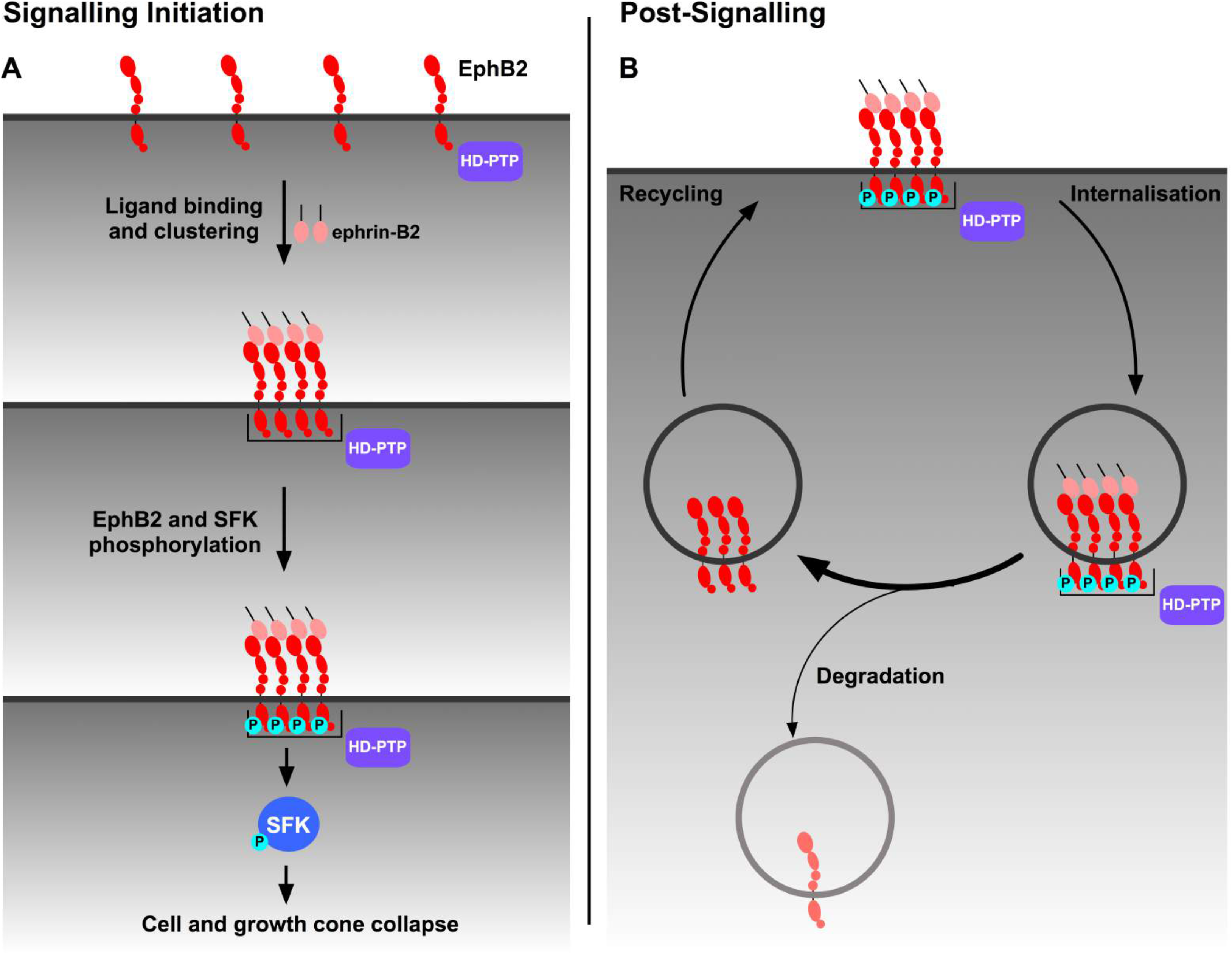
A model of the dual role of HD-PTP in EphB signalling. A) During signalling initiation, ephrin-B2 ligand multimers bind and cluster surface EphB2. The formation of the HD-PTP-EphB2 complex is facilitated by ephrin-B2 binding to EphB2. HD-PTP-depletion results in failure to form receptor clusters in response to ligand stimulation, suggesting HD-PTP promotes EphB2 multimerisation and/or ligand binding. This effect likely explains how HD-PTP loss can cause a failure to induce receptor phosphorylation, Src family kinase activation, as well as cytoskeleton destabilisation and cell collapse. B) Ligand-bound EphB2 complexes are ubiquitylated and eventually internalised in early endosomes, from where the receptor can be recycled back to the membrane or sorted to the endocytic pathway for lysosomal degradation. We propose that ligand-bound EphB2 complexes are protected by HD-PTP from degradation by promoting EphB2 recycling. HD-PTP’s deubiquitylase-recruiting function may play a role in this.

### ESCRT proteins in axon guidance

Our *in vivo* experiments uncover an important role of HD-PTP in nervous system development, through its function in the formation of connections between spinal motor neurons and their limb muscle targets. HD-PTP-deficient spinal motor axons, normally destined for the ventral limb nerve, enter the dorsal limb mesenchyme, which expresses ephrin-B2. Evidence that this is exerted through HD-PTP’s function in ephrin-B:EphB signalling includes the requirement of HD-PTP for ephrin-B2-induced motor neuron growth cone collapse *in vitro* and the requirement of ephrin-B2:EphB signalling for normal motor axon guidance *in vivo*. HD-PTP loss may also affect the response of spinal motor axon growth cones to other limb mesenchyme-derived signals important for motor axon guidance such as Netrin or Semaphorins (Poliak *et al.*, 2015; Huber *et al.*, 2005). However, cells and growth cones deficient in HD-PTP respond normally to Semaphorin3F and Semaphrin3A arguing against the involvement of HD-PTP in Semaphorin-mediated motor axon guidance. On the other hand, although there is no evidence of HD-PTP function in Netrin signalling, the ESCRT-II complex has been implicated in controlling the expression of DCC, a Netrin receptor (Konopacki *et al.*, 2016). Given the emerging prominence of post-translational control of axon guidance receptor function, the core ESCRT proteins could form a pervasive regulatory module, enabling endocytic processing of various axon guidance receptors, while ESCRT accessory proteins like HD-PTP may link such a module to specific axon guidance receptors.

### Conclusion

Our experiments linking ESCRT, a pervasive system controlling the fate of many transmembrane receptors, and Eph signalling, a rapid-action pathway underlying a wide variety of biological processes, bring many potential new insights into their understanding. For example, increased tumorigenesis caused by the loss of HD-PTP has been attributed to excessive surface receptor signalling (Gingras *et al.*, 2017), but in light of our data, could also be a consequence of impaired anti-cancer functions of Eph signalling (Pasquale, 2010). The control of Eph signalling by ESCRT proteins could thus be an important new therapeutic avenue in the context of tumorigenesis and other disorders involving Eph signalling, such as neurodegeneration (Cissé *et al.*, 2010; Van Hoecke *et al.*, 2012).

## Experimental procedures

### Animals

Fertilised chicken eggs (FERME GMS, Saint-Liboire, QC, Canada) were incubated at 38°C and staged according to Hamburger and Hamilton (1951).

### BioID and MS Data Analysis

BioID experiments were performed as described elsewhere, with modifications (Methot *et al.*, 2018). Briefly, Control and EphB2-OE Flp-In T-REx HEK293 cells (Invitrogen, Thermo Fisher Scientific, Waltham, MA) (all cell lines can be found in Supplementary Table 3) were cultured in 15 cm plates (Corning, New York) and treated with 1 μg/mL of tetracycline (Sigma Aldrich, St. Louis, MO) for 18 h. The following day, the medium was removed and cells were incubated in serum-free medium for 6 h in the presence of 50 μM biotin (Sigma Aldrich, St. Louis, MO) and pre-clustered ligand (Fc or eB2-Fc, 1.5 μg/mL, R&D Systems, Minneapolis, MN) or media. After 6 h of biotin and ligand treatment, the medium was removed, cells were scraped from the plates, washed 3 times with cold phosphate-buffered saline (PBS) in 15 mL tubes and cell pellets were stored at -80 °C. Cells were lysed in 1.5 mL radioimmunoprecipitation assay (RIPA) buffer and 1 μL of benzonase (MilliporeSigma, Burlington, MA) was added to each sample to degrade nucleic acids. Lysates were sonicated for 30 seconds (s) at 30% amplitude, in 10 s bursts with 2 s rest in between. Lysates were then centrifuged for 30’ at maximum speed at 4 °C. 70 μL of prewashed streptavidin beads (GE Healthcare Amersham, Little Chalfont, UK) were incubated with the remaining lysate for 3 h at 4 °C. Samples were spun down for 1’ at 2000 rpm at 4 °C and the supernatant was removed. Beads were re-suspended in 1.5 mL RIPA buffer and washed 3 times with RIPA buffer. Beads were then re-suspended in 1 mL of 50 mM Ammonium Bicarbonate (ABC, Bio Basic, Markham, Canada), washed 3 times with ABC and re-suspended in 100 μL of ABC. 1 μg of trypsin (Sigma Aldrich, St. Louis, MO) was added and samples were shaken at 37 °C for 16 h. The following day, samples were trypsin-digested for 2 h and spun down for 1’ at 2000 rpm at room temperature. Beads were washed 2 times in 100 μL of water (Caledon Laboratories, Georgetown, Canada) and combined with the collected supernatant. Formic acid (Sigma Aldrich, St. Louis, MO) was added to the supernatant for a final concentration of 5%. Samples were spun down for 10’ at maximum speed at room temperature, dried for 3 h at 30 °C (SpeedVac). Tryptic peptides were resuspended in 15 μL of 5% formic acid and stored at -80°C.

Peptides were analysed by high-pressure liquid chromatography (HPLC) coupled to Orbitrap Velos Mass Spectrometer (Thermo Fisher Scientific, Waltham, MA) at the IRCM proteomics core facility. Peptide search, identification of proteins and mass spectrometry (MS) data analysis were carried out as described elsewhere (Methot *et al.*, 2018). The BioID-MS data was analysed using ProHits (Liu *et al.*, 2010). Briefly, RAW files were converted to .mzXML using Proteowizard (Kessner *et al.*, 2008). Human RefSeq Version 57 and the iProphet tool integrated in ProHits (Shteynberg *et al.*, 2011) were used for peptide search and identification. Significance Analysis of INTeractome (SAINT) file inputs generated in ProHits were analysed through ProHits-viz (Knight *et al.*, 2017) to generate dot plots and to calculate WD-scores.

### Protein network analysis, clustering and functional annotation

Protein network and clustering analyses were generated via Cytoscape (Shannon *et al.*, 2003) as described elsewhere (Morris *et al.*, 2014), with modifications. Briefly, the BioID-MS data analysed in SAINT were imported to Cytoscape. Reviewed UniProtKB entries of the preys identified in SAINT were submitted into the ‘Enter Search Conditions’ text box and the existing protein-protein interaction data was imported from IntAct database (Hermjakob *et al.*, 2004). The BioID and public networks were merged by performing a union merge. Self-loops and duplicated edges were removed. MCL Cluster (Enright *et al.*, 2002) was used to visualise protein complexes and clusters. Functional annotation was performed using Gene Ontology (GO) terms. The known biological process or molecular function of prey proteins was analysed by using g:Profiler (Reimand *et al.*, 2016). Reviewed UniProtKB entries of the preys analysed in SAINT were submitted in the Query field on g:Profiler and the -log_10_ of corrected *p* values were used for GO enrichment and KEGG analysis.

### Biochemistry

For the co-immunoprecipitation assays, Control and EphB2-OE HEK293 cells were transfected with an HD-PTP-HA expression plasmid in 10 cm dishes using Lipofectamine 3000 (Invitrogen, Thermo Fisher Scientific, Waltham, MA), and one day later were incubated with DMEM (Gibco, Thermo Fisher Scientific, Waltham, MA) supplemented with 0.05% fetal bovine serum (Gibco, Thermo Fisher Scientific, Waltham, MA), 1% penicillin/streptomycin (100X, Gibco, Thermo Fisher Scientific, Waltham, MA) and 1 μg/mL of tetracycline (Sigma Aldrich, St. Louis, MO) for 18 h at 37 °C with 5% CO_2_. Control^shRNA^ and HD-PTP^shRNA^ HeLa cells (all cell lines can be found in Supplementary Table 3) were transfected with an EphB2-FLAG expression plasmid (gift from Dr. Matthew Dalva) using 2 mM calcium phosphate and stimulated 48 h after transfection. Stimulation with pre-clustered ligands (clustered with anti-Fc antibody for 30’ at room temperature) was for 15’ (Fc and eB2, 1.5 μg/mL) or 5’ (Fc and eB2, 1.0 μg/mL) at 37 °C. After a wash with PBS, cells were lysed with 1 M MgCl_2_, 2 M Tris-HCl pH 7.5, 3 M NaCl, 1% CHAPS (Bio Basic, Markham, Canada), 0.5 M sodium fluoride, 100 mM sodium orthovanadate and cOmplete proteinase inhibitor (25X, Roche, Basel, Switzerland). Lysates were spun down at 14,000 rpm for 15’ at 4 °C then supernatant was transferred and rotated with anti-FLAG beads (Sigma Aldrich, St. Louis, MO) for 3 h at 4 °C, washed 3 times with lysis buffer (same as above) and denatured with 6X Laemmli buffer (1:5).

For other biochemical assays, Control and EphB2 HeLa cells were incubated with DMEM supplemented with 0.05% fetal bovine serum, 1% penicillin/streptomycin and 1 μg/mL of tetracycline for 18 h at 37 °C with 5% CO_2_. Cells were lysed with 1 M Tris-HCl pH 8.0, 5 M NaCl, 1% NP-40 (Abcam, Cambridge, UK), phosSTOP (Sigma Aldrich, St. Louis, MO) and cOmplete proteinase inhibitor after a PBS wash. Lysates were spun down at 12,000 rpm for 5’ at 4 °C and denatured with 6X Laemmli buffer (1:5). Samples were run on 6–10% Bio-Tris polyacrylamide gels. Membranes (PVDF; Bio-Rad Laboratories, Hercules, CA) were activated with methanol (Sigma Aldrich, St. Louis, MO) for 2’ and put in either 1% BSA (Bio Basic, Markham, Canada), 0.05% Tween (Sigma Aldrich, St. Louis, MO) PBS or 5% milk blocking solution on a shaker for 45’ at room temperature and the following antibodies were applied: anti-GAPDH (1% BSA, 45’ at room temperature), anti-FLAG-HRP (1% BSA, 45’ at room temperature), anti-Streptavidin-HRP (1% BSA, 30’ at room temperature), anti-HA (5 % milk, 1 h at room temperature), anti-ß-actin (5% milk, 1 h at room temperature), and anti-HD-PTP (0.05% Tween PBS, overnight 4 °C). Information for all antibodies can be found in Supplementary Table 1. Membranes were activated with ECL (GE Healthcare Amersham, Little Chalfont, UK) and revealed with film (GE Healthcare Amersham, Little Chalfont, UK). Signal intensity and area of the immunoblot band was measured using ImageJ (NIH).

### Cell Culture

Control and EphB2-OE HEK293 and HeLa cells were generated by transfecting Flp-In T-REx HEK293 and Flp-In T-REx HeLa cells with either FLAG or EphB2-BirA*-FLAG expression plasmids using Lipofectamine 3000. Transfected cells were selected with hygromycin (200 μg/mL, Invitrogen, Thermo Fisher Scientific, Waltham, MA) for 15–16 days. Control^shRNA^ and HD-PTP^shRNA^ HeLa cells were generated by viral infection of either empty pLKO1 or HD-PTP shRNA pLKO1 (Sigma Aldrich, St. Louis, MO). After infection, cells were selected by puromycin (1 μg/mL, Gibco, Thermo Fisher Scientific, Waltham, MA) for 5–7 days and a western blot was performed to assess knock-down efficiency.

### HeLa Cell Collapse Assay

Control and EphB2 HeLa cells were seeded at 20,000 cells per coverslip (VWR, Avantor, Center Valley, PA). After 24 hours, cells were incubated in DMEM supplemented with 0.05% fetal bovine serum, 1% penicillin/streptomycin and 1 μg/mL of tetracycline for 18 h at 37 °C with 5% CO_2_ and stimulated with pre-clustered eB2 or Fc the next day. Control^shRNA^ and HD-PTP^shRNA^ HeLa cells were cultured in DMEM supplemented with 10% fetal bovine serum, 1% penicillin/streptomycin and puromycin (1 μg/mL) at 37 °C with 5% CO_2_. These cells were transfected in 6-well plates (Sarstedt, Thermo Fisher Scientific, Waltham, MA) using Lipofectamine 3000, seeded at 20,000 cells per coverslip and stimulated 48 h after transfection and 24 h after being seeded.

### Chick *in ovo* electroporation and CRISPR guides

Chicken spinal cord electroporation of expression plasmids was performed at HH st. 18/19 as described (Croteau & Kania, 2011). Guide RNAs were designed against the HD-PTP *Gallus gallus* genomic sequence using CHOPCHOP (Labun *et al.*, 2016) and were verified for specificity using the NCBI BLAST tool (Gish & States, 1993). The pX330 plasmid (#42230 obtained from Addgene) was modified by subcloning T2A-EGFP cassette downstream and in frame to Cas9, producing pX3361. Guide RNA oligos (Synthego, Menlo Park, CA) were synthetically made and cloned in the pX3361 plasmid. Guide RNA sequences are available upon request.

### *In situ* mRNA localisation and immunohistochemistry

*In situ* mRNA detection and immunofluorescence were performed as described (Kao & Kania, 2011) or using standard methods. Probe sequences are available upon request.

For non-permeabilised assays on LMC growth cones, tissue was exposed to ligands for 15’ and placed on ice, and a 5’ blocking step was performed by replacing half the media with PBS containing 2% BSA (final, 1% on tissue) and incubating at 4 °C. Half of the media was then replaced with motor neuron media (Neurobasal media (Gibco, Thermo Fisher Scientific, Waltham, MA) supplemented with B-27 (1:50, Gibco, Thermo Fisher Scientific, Waltham, MA), 0.5 mM L-Glutamate (Sigma Aldrich, St. Louis, MO), 25 mM L-Glutamine (Gibco, Thermo Fisher Scientific, Waltham, MA), and 1% penicillin-streptomycin containing primary antibodies against EphB2 (1 in 1000) and EEA1 (1 in 500) as control, and incubated for 30’ at 4 °C (all antibody details can be found in Supplementary Table 1). Tissue was then fixed with a mixture of 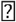 30% sucrose (Bio Basic, Markham, Canada) and 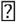 4% PFA (Sigma Aldrich, St. Louis, MO) for 15’ at 4 °C. Three washes with PBS were followed by adding secondary antibodies (final, 1 in 1000 in PBS) for 1 h at 4 °C. Finally, three quick washes with PBS were followed by mounting in Mowiol (Sigma Aldrich, St. Louis, MO). For the permeabilised control, fixation occurred after ligand incubation and before primary antibody staining, primaries were added in media with added Triton X-100 (0.3%, Sigma Aldrich, St. Louis, MO), and secondary antibodies in PBS with added Triton X-100 (0.3%). Otherwise, all concentrations, incubation times, and temperatures were identical.

For non-permeabilised assays on HeLa cells, cells were exposed to ligands for 5’ and placed on ice immediately. Blocking was done by replacing half the media with PBS containing 2% BSA (final, 1% on tissue) for 5’ at 4 °C, whereupon half the media was replaced with DMEM supplemented with 0.05% fetal bovine serum, 1% penicillin/streptomycin, and containing primary antibodies against EphB2 (1 in 1000) and EEA1 (1 in 500) as control, and incubated for 30’ at 4 °C. Secondary staining was performed as in growth cones.

### Motor Neuron Culture

HH st. 25 chick embryos were harvested and dissected to isolate the motor column of the spinal cord. Tissue was dissociated with 0.25% trypsin (Life Technologies) in Ca^2+^/Mg^2+^ Hanks’s Solution (Invitrogen, Thermo Fisher Scientific, Waltham, MA) deactivated by 1M MgSO4 (Invitrogen, Thermo Fisher Scientific, Waltham, MA) and 12500 U/mL DNAse (Worthington Industries, Columbus, OH). Cells were spun down at 1000 rpm for 5’ at room temperature, and resuspended in Neurobasal media supplemented with 1% fetal bovine serum, 0.01% Glutamax (Invitrogen, Thermo Fisher Scientific, Waltham, MA), and 0.01% penicillin/streptomycin then titrated. 20,000 cells were seeded onto laminin-coated (20 μg/mL; Invitrogen, Thermo Fisher Scientific, Waltham, MA) coverslips and incubated at 37 °C with 5% CO_2_. Cells were stimulated with pre-clustered eB2 or Fc one day after being seeded.

### Microscopy and Image Quantification

High magnification images were taken using ZEN 2010 on a Zeiss LSM 700 confocal microscope. Lower magnification pictures were taken using LasX on a Leica DFC 488 light microscope. *In situ* hybridization images were taking using OsteoMeasure on a Leica DM 4000 light microscope. Axon projection, mean intensity of signal, cell area, motor neuron numbers of limb sections were quantified using ImageJ (NIH) and methods previously described (Kao *et al.*, 2009).

### PCR

Chick HH st. 25 spinal cords were digested with 100 μg/mL proteinase K (Thermo Fisher Scientific, Waltham, MA) in SDS buffer (100 mM Tris pH 8.5, 5 mM EDTA, 200 mM NaCl, 0.2% SDS) at 55 °C for 3 h. The extracted DNA was precipitated by isopropanol, washed with 70% ethanol, and re-suspended in ddH_2_O. PCR amplification of HD-PTP genomic locus in control or HD-PTP^CRISPR^-electroporated tissue was performed using Qiagen Master Mix and the following primers: forward outside primer (tttggggcagacagacatct), reverse outside primer (tatctttcgcacccctgctc). Nested PCR was performed with 1 μL of the previous PCR reaction product, with Qiagen Master Mix and the following primers: forward inside (agaaaggcacctgctccca) primer, reverse inside primer (ttccagtcacacagcagctg). PCR products were then visualised on a 1% agarose gel after electrophoresis.

### Pulse Chase

Control^shRNA^ and HD-PTP^shRNA^ HeLa cells were transfected in 10 cm dishes (as above) with an EphB2-FLAG expression plasmid (gift from Dr. Matthew Dalva) using Lipofectamine 3000. After 24 h, 80,000 cells were seeded into 24 well plates and stimulated with pre-clustered eB2 or Fc 24 h later. Cells were pulsed with 10 μg/mL cycloheximide and 1μg/mL of pre-clustered ligand (Fc or eB2, 1.0 μg/mL). Samples were collected at various time points and total EphB2 protein quantity was analysed by immunoblotting.

### Statistical Analysis

Data from the experimental replicates were evaluated using Prism (GraphPad Software, California). Means of individual experiments were compared and underwent various statistics. For 3 or more conditions, one-way ANOVA was used, followed, if necessary, by Student’s *t-*tests corrected for multiple comparisons. For comparing 2 conditions with less than four replicates, we assumed normal distributions and analysed them with Student’s *t-*tests. For the growth cone collapse assay, which entailed categorical analysis, Fisher’s exact test was used. The threshold for statistical significance was set at 0.05.

## Acknowledgements

The authors thank J. Cardin and M. Liang for technical assistance, L. Delorme for secretarial assistance, and N. Bisson for critical comments on an earlier version of the manuscript. D.M. was a recipient of a Mexican National Council for Science and Technology (CONACYT) international PhD scholarship, received funding from the McGill University Integrated Program in Neuroscience, and is currently funded by a Swiss Government Excellence Postdoctoral Scholarship. H.B. was supported by a doctoral training award from the Fonds de recherché du Québec – Santé (FRQS) (#33603). G.D. was supported by a postdoctoral training award from the FRQS. This work was supported by grants from the Canadian Institutes of Health Research (PJT-152966 to A.P, MOP-97758 and MOP-77556 to A.K.), the Canadian Cancer Society Research Institute (705376 to A.P.), NSERC (RGPIN-2016-04808 to J.F.C.), and Brain Canada, Canadian Foundation for Innovation, and the W. Garfield Weston Foundation to A.K. J.F.C. holds the TRANSAT chair in breast cancer research. A.K. and J.F.C. were also supported by FRSQ Chercheur-boursier Senior career awards.

## Supplemental Figure Legends

**Figure S3.**
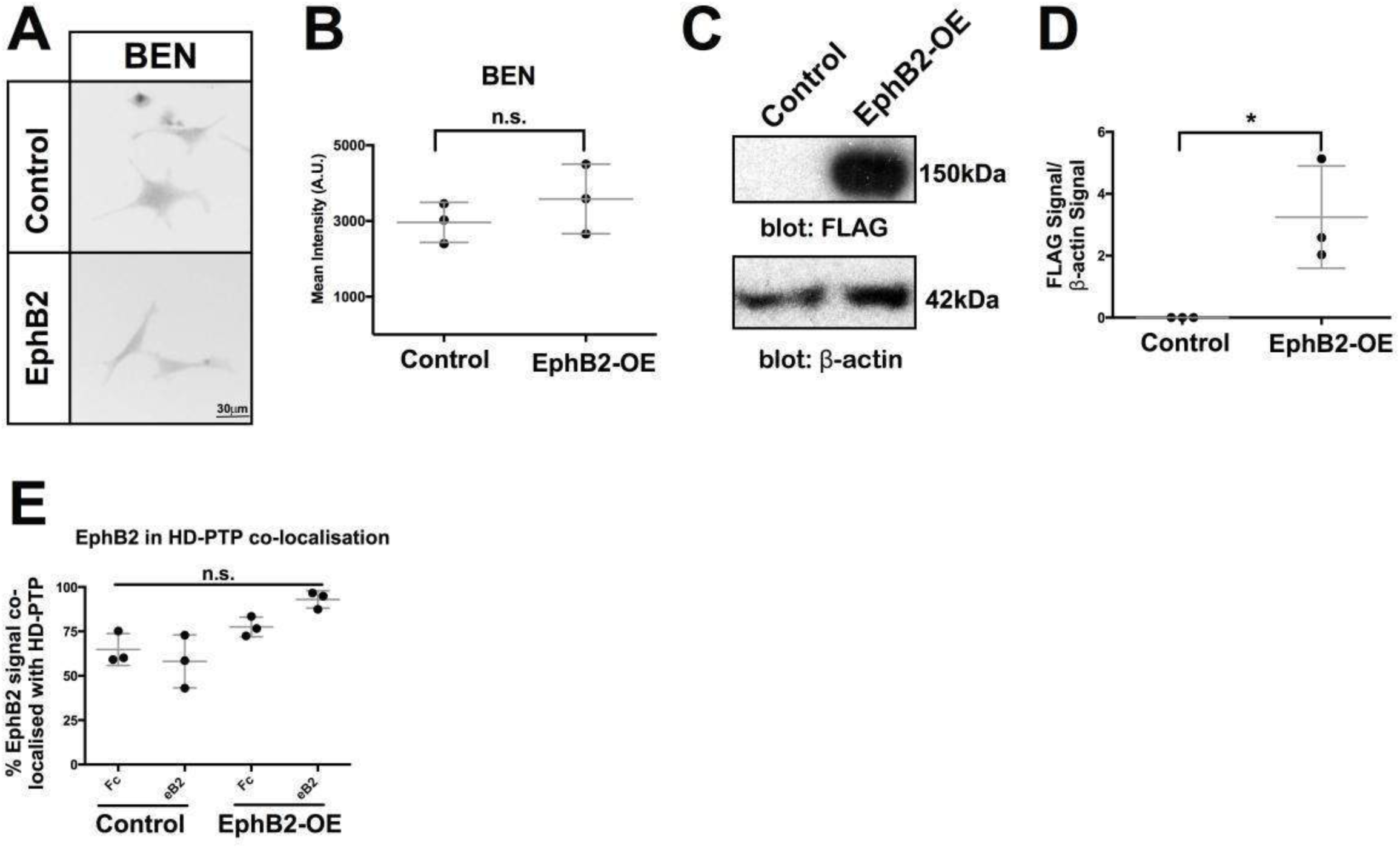
HD-PTP and EphB2 expression and localisation are linked in HeLa cells. A) Representative inverted grayscale fluorescent images of Control HeLa and EphB2-OE HeLa immunostained with anti-BEN antibody. B) Quantification of BEN mean pixel intensity in HeLa cells (*n* = 3, 60–80 cells/*n*; Student’s *t*-test). C) Western blot of FLAG showing EphB2-BirA*-FLAG at 150 kDa and FLAG, not shown in blot, at 5 kDa. D) Quantification of FLAG signal relative to β-actin signal (*n* = 3, Student’s *t*-test). **E)** Quantification of EphB2 localisation in HD-PTP-positive domains. EphB2 is found in HD-PTP-containing puncta in equal proportions in Control and EphB2-OE HeLa cells (*n* = 3, 10–12 cells/*n*; Student’s *t*-test). Values are plotted as mean ± SD. All values can be found in Supplementary Table 4. kDa: kilodalton; eB2: ephrin-B2-Fc; * *p* < 0.05; n.s., not significant. Scale bar in A: 30 μm.

**Figure S4.**
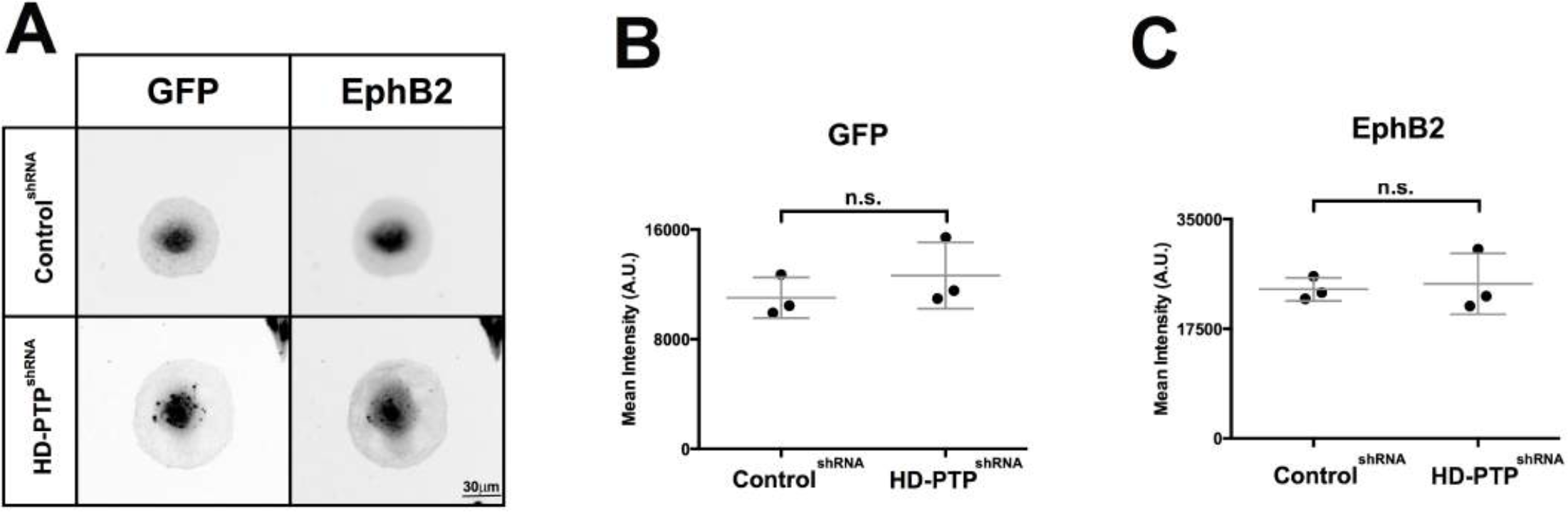
HD-PTP is required for ephrin-B2-induced HeLa cell collapse. A) Representative inverted grayscale fluorescent images of Control^shRNA^ and HD-PTP^shRNA^ HeLa cells transfected with EphB2-GFP plasmid, showing GFP and anti-EphB2 signals. B) Quantification of GFP mean pixel intensity in Control^shRNA^ and HD-PTP^shRNA^ HeLa cells transfected with EphB2-GFP plasmid (*n* = 3, 60–80 cells/*n*; Student’s *t*-test). C) Quantification of EphB2 mean pixel intensity in Control^shRNA^ and HD-PTP^shRNA^ HeLa cells transfected with EphB2-GFP plasmid (*n* = 3, 60–80 cells/*n*; Student’s *t*-test). Values are plotted as mean ± SD. All values can be found in Supplementary Table 4. n.s.: not significant. Scale bar in A: 30 μm.

**Figure S5.**
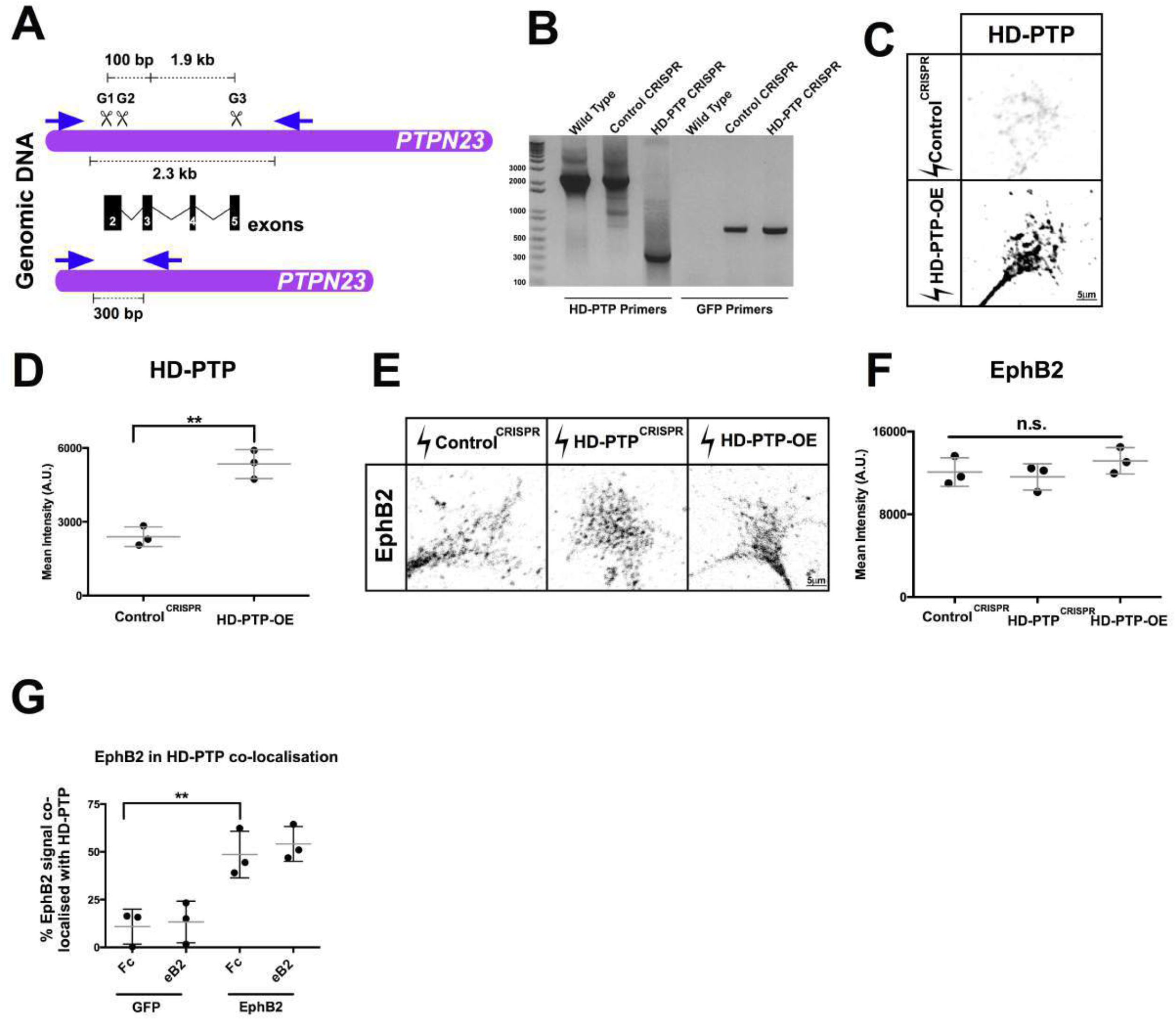
HD-PTP CRISPR knockdown strategy. A) Schematic depicting the *PTPN23* (chicken gene encoding HD-PTP) genomic locus, the location CRISPR guides G1, G2 and G3 and PCR primers (arrows). The three guide RNAs produce deletions of exons 2–5. B) Representative genomic PCR using the HD-PTP primers in (A) and GFP primers in DNA from a wild-type chick spinal cord, a Control^CRISPR^-electroporated spinal cord, and a HD-PTP^CRISPR^-electroporated spinal cord. HD-PTP primers show a full-length 2300 bp band in wild-type spinal cord and Control^CRISPR^ spinal cord, and a cleaved 300 bp band in the HD-PTP^CRISPR^ spinal cord. GFP primers show no band in wild-type spinal cords, and a 750 bp band in both Control^CRISPR^ and HD-PTP^CRISPR^ spinal cords (*n* = 3). C) Representative images of cultured Control^CRISPR^ and HD-PTP-OE chick HH St. ## LMC growth cones stained with the anti-HD-PTP antibody. D) Quantification of HD-PTP mean pixel intensity of Control^CRISPR^ and HD-PTP-OE growth cones (*n* = 3, 10–12 growth cones/*n*; Student’s *t*-test). E) Representative images of growth cones of LMC neurons electroporated either with Control^CRISPR^, HD-PTP^CRISPR^ or hHD-PTP-FLAG plasmids, stained with anti-EphB2 antibody. F) Quantification of EphB2 mean pixel intensity in LMC growth cones electroporated with Control^CRISPR^, HD-PTP^CRISPR^ or hHD-PTP-FLAG (*n* = 3, 10–12 growth cones/*n*; Student’s *t*-test). G) Quantification of EphB2 localisation in HD-PTP-positive puncta in electroporated growth cones. Approximately 50% of EphB2 is found in HD-PTP-containing sites when EphB2-GFP is electroporated, compared to 20% in GFP-electroporated LMC growth cones (*n* = 3, 10–12 growth cones/*n*; Student’s *t*-test). Values are plotted as mean ± SD. All values can be found in Supplementary Table 4. G: guide RNA; bp: base pairs; kb: kilobase; ** *p* < 0.01; n.s., not significant. Scale bars: C and E) 5 μm. Inverted grayscale fluorescent images.

**Figure S6.**
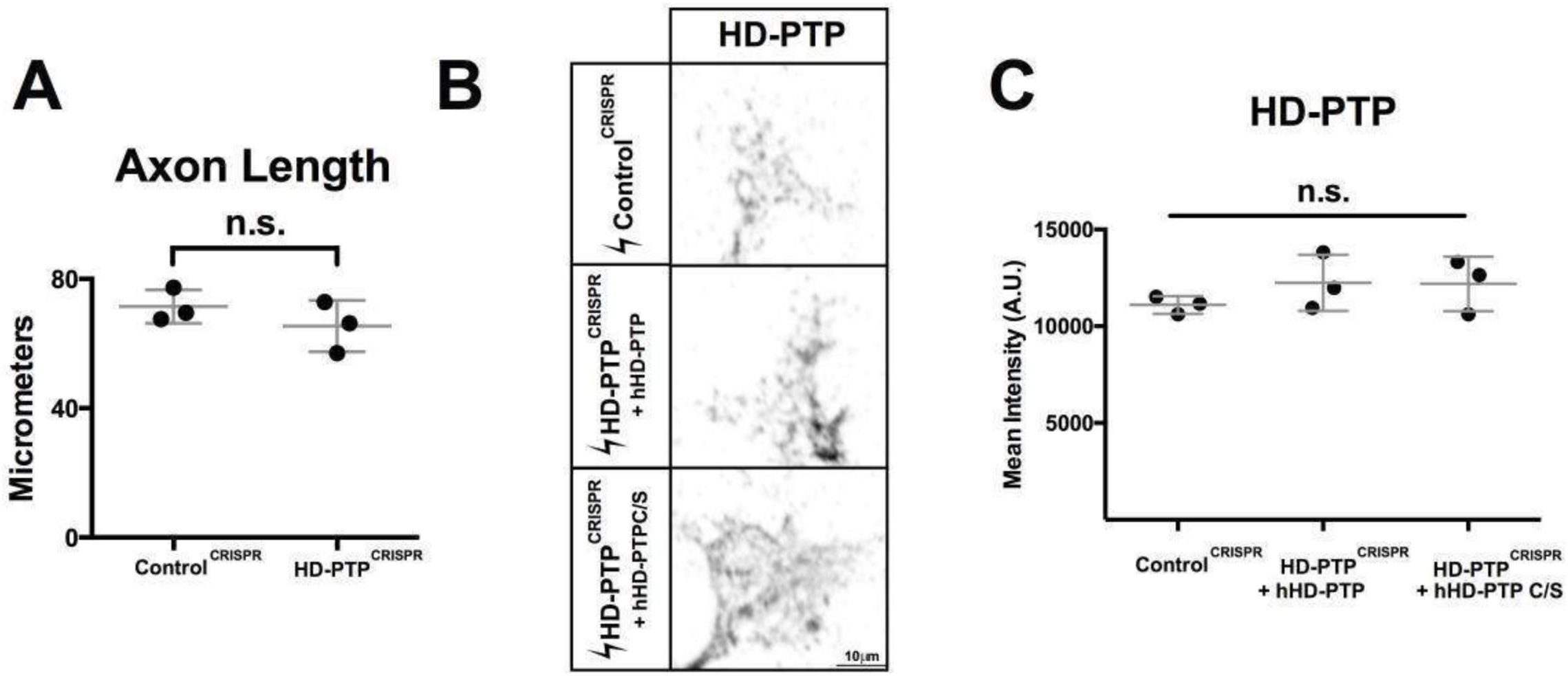
Medial LMC growth cones require HD-PTP for ephrin-B2-induced collapse. A) Quantification of Control^CRISPR^ and HD-PTP^CRISPR^-electroporated LMC motor neuron axon length *in vitro* (*n* = 3, 30–50 axons/*n*; Student’s *t*-test). B) Representative inverted grayscale fluorescent images of Control^CRISPR^, HD-PTP^CRISPR^ + hHD-PTP, and HD-PTP^CRISPR^ + hHD-PTP C/S LMCm growth cones stained with the anti-HD-PTP antibody. C) Quantification of HD-PTP mean pixel intensity of Control^CRISPR^, HD-PTP^CRISPR^ + hHD-PTP, and HD-PTP^CRISPR^ + hHD-PTP C/S LMCm growth cones (*n* = 3, 10–12 growth cones/*n*; one-way ANOVA). Values are plotted as mean ± SD. All values can be found in Supplementary Table 4. n.s., not significant. Scale bar in B: 10 μm.

**Figure S8.**
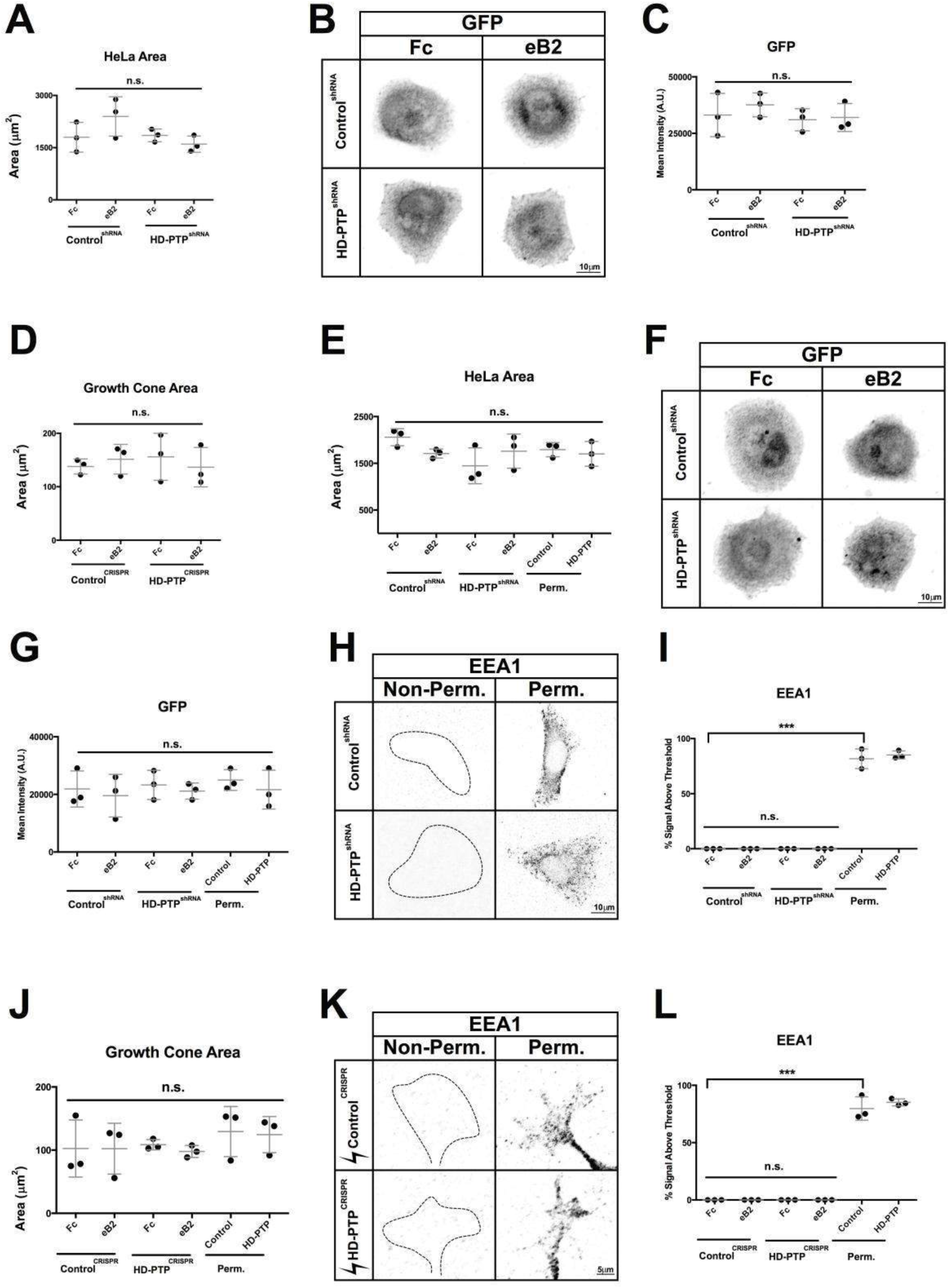
HD-PTP is required for ligand-induced EphB2 phosphorylation, SFK phosphorylation, and EphB2 surface patching. A) Cell area quantification in Control^shRNA^ and HD-PTP^shRNA^ HeLa cells transfected with the EphB2-GFP plasmid and incubated for 5’ with 1 μg/mL eB2 or Fc as controls for the anti-phospho-Y418-SFK experiment in Fig. 8C (*n* = 3, 10–12 cells/*n*; one-way ANOVA). B) Representative images of Control^shRNA^ and HD-PTP^shRNA^ HeLa cells transfected with the EphB2-GFP plasmid used in anti-phospho-Y418-SFK experiment in Fig. 8C. C) GFP signal quantification in Control^shRNA^ and HD-PTP^shRNA^ HeLa cells transfected with the EphB2-GFP plasmid and incubated for 5’ with 1 μg/mL eB2 or Fc as controls for the anti-phospho-Y418-SFK experiment in Fig. 8C (*n* = 3, 10–12 cells/*n*; one-way ANOVA). D) Area quantification in Control^CRISPR^ and HD-PTP^CRISPR^ LMC growth cones that were incubated for 15’ with 10 μg/mL eB2 or Fc, as controls for anti-phospho-Y418-SFK experiment in Fig. 8E (*n* = 3, 10–12 cells/*n*; one-way ANOVA). E) Cell area quantification in Control^shRNA^ and HD-PTP^shRNA^ HeLa cells transfected with the EphB2-GFP plasmid and incubated for 5’ with 1 μg/mL eB2 or Fc as controls for the non-permeabilised vs. permeabilised EphB2 experiment in Fig. 8G (*n* = 3, 10–12 cells/*n*; one-way ANOVA). F) Representative images of HD-PTP^shRNA^ and Control^shRNA^ HeLa cells transfected with the EphB2-GFP plasmid as controls for the non-permeabilised vs. permeabilised EphB2 experiment in Fig. 8G. G) GFP signal quantification in Control^shRNA^ and HD-PTP^shRNA^ HeLa cells transfected with the EphB2-GFP plasmid and incubated for 5’ with 1 μg/mL eB2 or Fc as controls for the non-permeabilised vs. permeabilised EphB2 experiment in Fig. 8G (*n* = 3, 10–12 cells/*n*; one-way ANOVA). H) Representative images of Control^shRNA^ and HD-PTP^shRNA^ HeLa cells, non-permeabilised vs. permeabilised, stained with the anti-EEA1 antibody. I) EEA1 signal quantification in Control^shRNA^ and HD-PTP^shRNA^ HeLa cells transfected with EphB2-GFP plasmid and incubated for 5’ with 1 μg/mL eB2 or Fc as controls for the non-permeabilised vs. permeabilised EphB2 experiment in Fig. 8G (*n* = 3, 10–12 cells/*n*; one-way ANOVA). J) Area quantification in Control^CRISPR^ and HD-PTP^CRISPR^ LMC growth cones that were incubated for 15’ with 10 μg/mL eB2 or Fc as controls for the non-permeabilised vs. permeabilised EphB2 experiment in Fig. 8I (*n* = 3, 10–12 growth cones/*n*; one-way ANOVA). K) Representative images of Control^CRISPR^ and HD-PTP^CRISPR^ LMC growth cones, non-permeabilised vs. permeabilised, stained with an anti-EEA1 antobody. L) EEA1 signal quantification in Control^CRISPR^ and HD-PTP^CRISPR^ LMC growth cones that were incubated for 15’ with 10 μg/mL eB2 or Fc as controls for the non-permeabilised vs. permeabilised EphB2 experiment in Fig. 8I (*n* = 3, 10–12 growth cones/*n*; one-way ANOVA). Values are plotted as mean ± SD. All values can be found in Supplementary Table 4. eB2: ephrin-B2-Fc; *** *p* < 0.001; n.s.: not significant. Scale bars: B, F, H) 10 μm, K) 5 μm. Inverted grayscale fluorescent images.

## Supplementary Tables

**Supplemenatary Table 1:** Antibodies and reagents

| **Antigen/**<br>**recombinant protein** | **Source Species** | **Dilution /**<br>**concentration** | **Source/reference** |
| --- | --- | --- | --- |
| Foxp1 | Rabbit | 1:1000 | Abcam |
| Isl1 | Mouse | 1:100 | DSHB |
| GFP | Rabbit | 1:1000 | Invitrogen |
| Ephrin-B2-Fc | Mouse | CoIP = 1.5 µg/mL<br>HeLa = 1.0 µg/mL<br>Growth Cones = 10<br>µg/mL | R&D systems |
| Sema3A-Fc | Human | 300 ng/mL | R&D Systems |
| Sema3F-Fc | Mouse | 300 ng/mL | R&D Systems |
| Fc | Human | Matched with ephrin-B2 and Sema<br>concentrations. | R&D Systems |
| EphB2 | Goat | 1:1000 | R&D Systems |
| EEA1 | Rabbit | 1:1000 | Abcam |
| Anti-Fc | Goat | 1:4 mass ratio to<br>ephrin-B2 | Sigma Aldrich |
| Anti-Fc | Mouse | 1:4 mass ratio to<br>ephrin-B2 | Sigma Aldrich |
| Tuj1 | Mouse | 1:1000 | Covance |
| 568-Phalloidin |  | 1:500 | Life Technologies |
| HA | Mouse | 1:2000 | Sigma Aldrich |
| Flag | Mouse | 1:200 | Sigma Aldrich |
| Flag-HRP | Mouse | 1:8000 | Sigma Aldrich |
| Beta-actin | Mouse | 1:5000 | Sigma Aldrich |
| HD-PTP | Rabbit | 1:2000 (WB)<br>1:200 (IF) | (Gingras et al., 2009) |
| Phosphotyrosine Y20 | Mouse | 1:2000 | BD Biosciences |
| pSFK-Y418 | Rabbit | 1:500 | Life Technologies |
| Streptavidin-HRP | Mouse | 1:25000 | Sigma Aldrich |
| GAPDH-HRP | Mouse | 1:2000 | Sigma Aldrich |

**Supplemenatary Table 2:** Plasmids

| **Plasmid** | **Species** | **Backbone** |
| --- | --- | --- |
| EphA4 CRISPR | targeting Chick | pX3361 |
| EphB2-GFP | Mouse | pN2-GFP |
| EphB2-FLAG | Mouse | pCMV |
| GFP | *Aequorea victoria* | pN2-GFP |
| HD-PTP CRISPR | targeting Chick (3 guides) | pX3361 |
| HD-PTP-FLAG | Human | pcDNA3 |
| HD-PTP(C/S)-FLAG | Human | pcDNA3 |
| HD-PTP-HA | Human | pcDNA3 |

**Supplemenatary Table 3:**
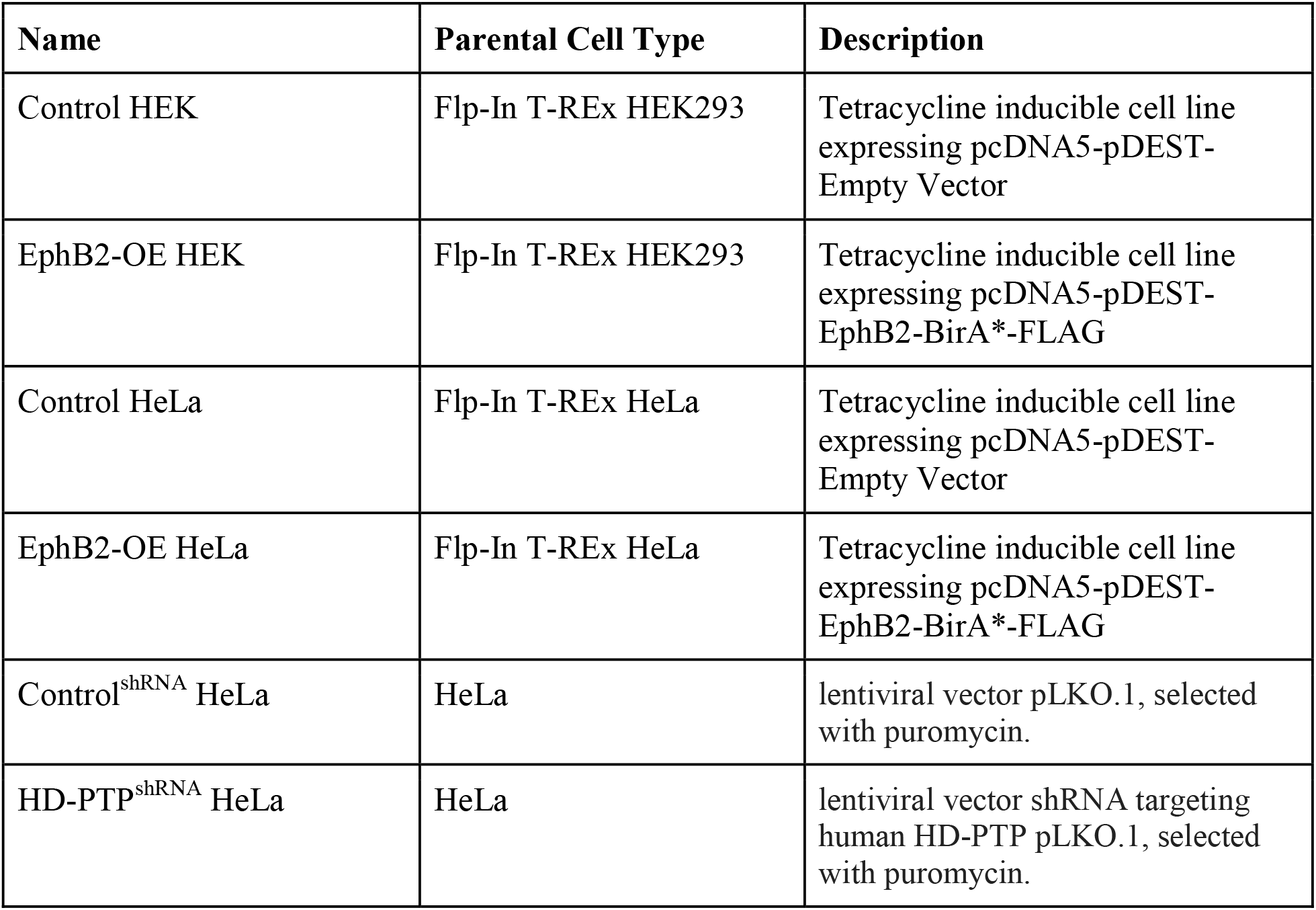
Cell lines

**Supplemenatary Table 4:**
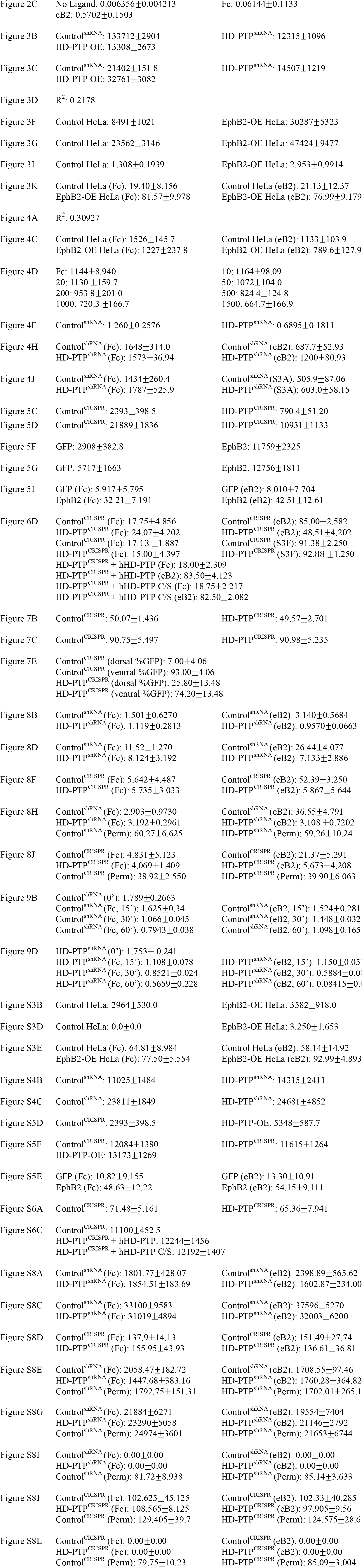
Quantifications of main & supplemental figures (All values are expressed as mean±SD

**Supplemenatary Table 5:**
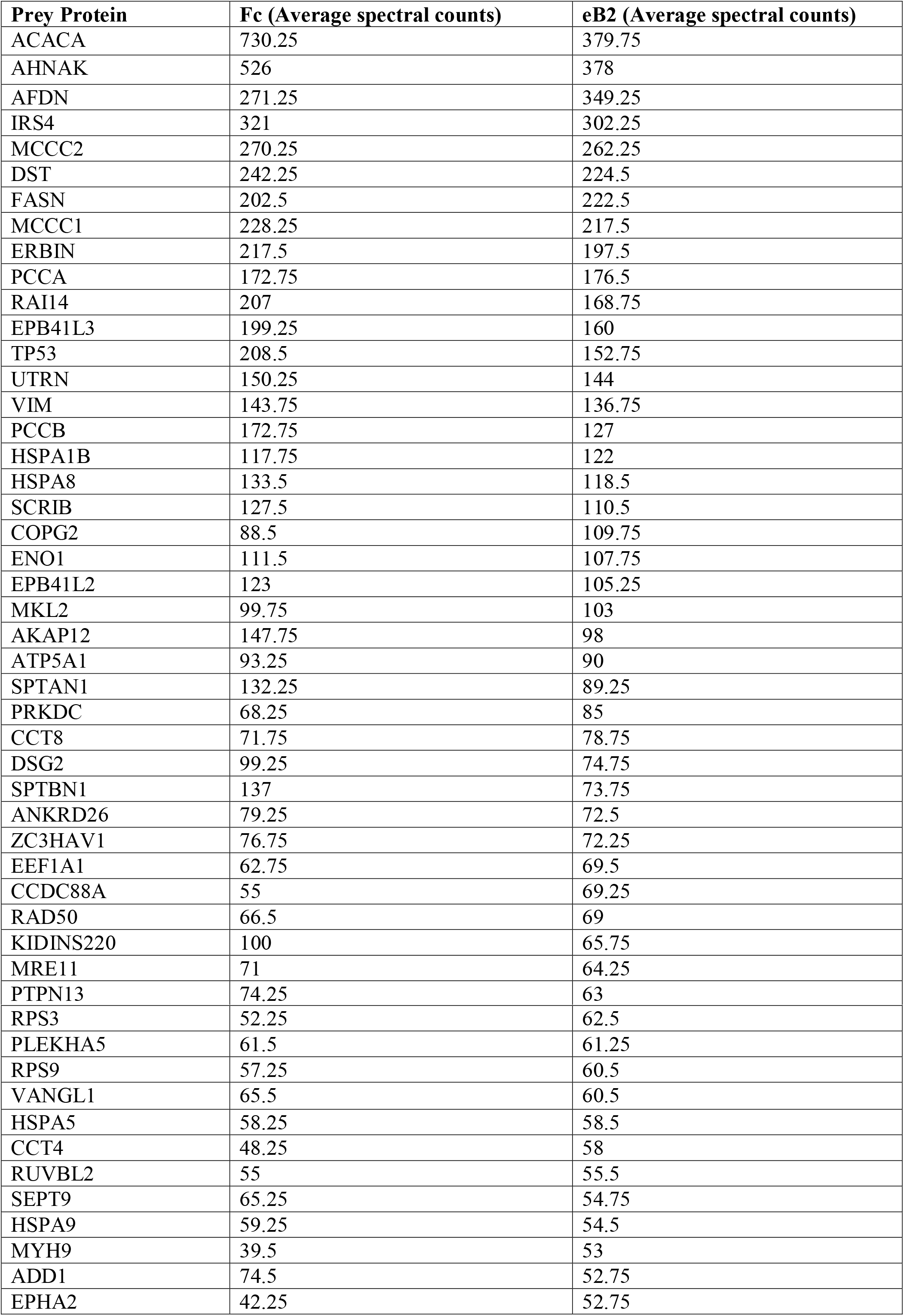
Top 50 BioID hits. Highest average spectral counts (*n* = 4) in eB2 stimulated condition after filtering with BirA*-FLAG-EGFP and empty vector MS datasets (Lambert et al., 2015).

## Notes

**Competing interests** The authors declare no financial or non-financial competing interests.

